# A NOVEL UNBIASED SEED-BASED RNAi SCREEN IDENTIFIES SMALL RNAs THAT INHIBIT ANDROGEN SIGNALING AND PROSTATE CANCER CELL GROWTH

**DOI:** 10.1101/2022.09.29.510140

**Authors:** Joshua M Corbin, Constantin Georgescu, Lin Wang, Jonathan D Wren, Magdalena Bieniasz, Chao Xu, Adam S Asch, Maria J Ruiz-Echevarría

## Abstract

Blocking androgen receptor signaling is the mainstay of therapy for advanced prostate cancer (PCa). However, acquired resistance to single agents targeting this pathway results in the development of lethal castration resistant PCa. Combination therapy approaches represent a promising strategy for the treatment of advanced disease. Here we explore a therapeutic strategy for PCa based on the ability of sh/siRNAs to function essentially as miRNAs and, via seed sequence complementarity, induce RNA interference of numerous targets simultaneously. We developed a library that contained shRNAs with all possible seed sequence combinations to identify those ones that most potently reduce cell growth and viability when expressed in PCa cells. Validation of some of these RNAi sequences indicated that the toxic effect is associated with seed sequence complementarity to the 3’-UTR of AR coregulatory and essential genes. In fact, expression of siRNAs containing the identified toxic seed sequences led to global inhibition of AR-mediated gene expression and reduced expression of cell cycle genes. When tested in mice, the toxic shRNAs also inhibited castration resistant PCa and exhibited therapeutic efficacy in pre-established tumors. This multi-targeted RNAi approach may be a promising therapeutic strategy for PCa.

## INTRODUCTION

Prostate cancer (PCa) is the second leading cause of cancer related death in men, and a major public health concern.^1^ The high mortality rate is largely due to development of resistance to current therapies for advanced disease. Androgen deprivation therapy (ADT), which prevents androgen stimulation, exploits the dependency of PCa cells on androgen signaling for growth and survival, and it is the standard of care for metastatic disease.^2^ However, while ADT is initially effective, most patients relapse with castration-resistant PCa (CRPC), a lethal form of the disease in which the PCa cells acquire the ability to grow in the androgen depleted environment while remaining dependent on androgen receptor (AR) activity.^3,4^ This reliance has led to the development of second-generation antiandrogens, or androgen biosynthesis blockers, which show effective but short-lived responses, leading to secondary resistance, and even to complete AR independence.^5,6^

Critical to AR-mediated signaling is the recruitment of AR coregulators which modulate and specify its transcriptional response, inferring an important role for AR coregulators in PCa. Accordingly, mechanisms of resistance to hormonal therapies often involve not only genetic and epigenetic alterations in the *AR* itself (i.e. amplification, mutations and splice variants), but also changes in the expression of AR coregulators.^3,4^ In fact, increased expression of AR coregulators generally correlates with aggressive disease and poor clinical outcome,^7^ suggesting that targeting AR coregulators, or their interaction with the AR, are plausible therapeutic alternatives to block androgen signaling. However, our understanding of the composition and the function of the AR coregulator complexes is incomplete, and targeting individual AR coregulators has been largely unsuccessful. This is likely due to functional redundancies and compensatory mechanisms, and points to the need of finding therapeutic approaches that target multiple AR coregulators.

RNA interference (RNAi) may provide such an approach. Small non-coding RNAs (miRNAs, shRNAs, siRNAs), bind and recruit the RNA induced silencing complex (RISC) to the 3’UTR of its mRNA targets through seed sequence (nucleotides 2-7 or 8 of the guide strand of the mi/si/shRNA) complementarity, resulting in mRNA degradation and/or translation inhibition.^8–14^ Small RNAs are predominantly implicated in regulating critical biological pathways and it has been proposed that the targets of a single miRNA are generally functionally associated (networks).^15,16^ Recently published studies from our lab identified sh/siRNAs that reduce the expression of AR coregulatory and essential gene networks in PCa cells through a seed-mediated mechanism, resulting in androgen signaling inhibition and cell death.^17^ Together with other studies demonstrating the ability of RNAi to target essential gene networks to induce cell death in other cancer cell types,^18–20^ our work provides a rationale for pursuing RNAi of AR signaling networks as a potential therapeutic strategy for PCa.

In this study, we developed a novel unbiased, seed-based, shRNA negative selection screen to identify shRNAs that most effectively inhibit PCa cell growth (toxic). Our results indicate that the toxic shRNAs inhibit global androgen signaling and suggest a clear link between the effect on PCa cell growth and viability and RNAi targeting of AR coregulatory and essential gene networks. Importantly, our results validate the therapeutic potential of the toxic shRNAs *in vivo,* and point to a therapeutic advantage with respect to individual coregulator targeting in PCa cells. Overall, our results suggest that RNAi-mediated targeting of androgen signaling regulatory and essential gene networks might present a promising therapeutic strategy for the treatment of PCa.

## RESULTS

### A novel unbiased RNAi screen for the Identification of negatively selected shRNAs in PCa cells

In order to identify novel RNAi sequences that potently inhibit PCa cell growth, we conducted a novel unbiased negative selection seed-based shRNA screen in PCa cell lines. We used a lentiviral based library expressing 15,572 unique shRNAs from a doxycycline (Dox) inducible promoter, and that differ in the 7-nucleotide seed sequence (nucleotides 2-8 of the guide strand). All possible seven nucleotide permutations within the guide strand 7mer seed sequence were included, while the sequence context surrounding the seed was identical for all shRNAs (Figure 1A). We selected an androgen dependent PCa cell line (LNCaP) and three CRPC cell lines (LNCaP abl, LNCaP-95 and 22Rv1) for the screen. 22Rv1 cells were grown in the presence and absence of 10 nM dihydrotestosterone (DHT) due to different AR regulated signatures, and their ability to grow long-term, in both conditions. 22Rv1 and LNCaP-95 cells express the constitutively active AR splice isoform, AR-V7. Cells were transduced with the lentiviral library, and after selection a sample was taken for a day 0 starting distribution while the rest were grown in the presence of doxycycline for ten doublings to allow for negative selection prior to NGS for quantification and identification of the depleted sequences. Z-scores were calculated as previously described,^21^ and 88 depleted shRNAs were selected according to our criteria (Figures 1B and S1, S2). The 7mer seeds from depleted shRNAs had a tendency towards being G/T rich (Figure 1C, 1D, S3). We identified eleven 6mer seed sequences (nucleotides 2-7 of the guide strand) that were present within multiple depleted shRNAs (Figures 1A and 1D); a finding that is of significance, as seed sequences can function as 6 or 7mers.^10–14^ These seeds include that of the miR-34 tumor suppressor miRNA^19,22,23^ (GGCAGT) which has been found to target the AR in PCa cell lines,^24^ highlighting the viability of the screen to identify shRNAs that target genes that are essential for PCa growth.

**Figure 1:**
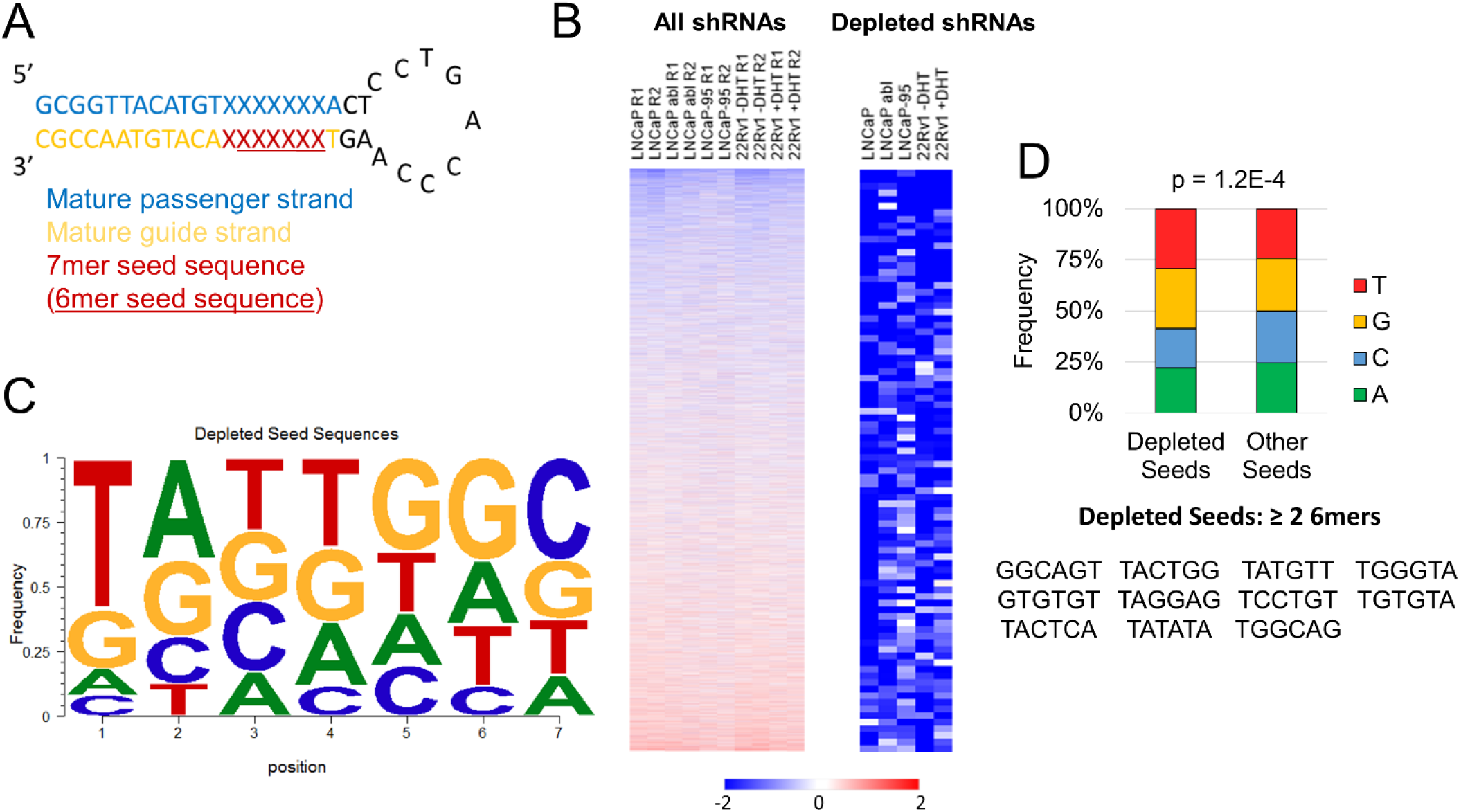
Identification of negatively selected shRNA sequences. **A)** Schematic of the shRNA construct used for the screen. **B)** Heatmaps showing Z-scores of the shRNAs (n=15,572) after 10 doublings (Left). Average Z-Scores of selected depleted shRNAs (n=88) are shown (Right). Red or blue indicates that the corresponding shRNA was positively or negatively selected respectively. **C)**. Motif stack plot showing the relative frequency of each nucleotide at each position for the 7mer seed sequences of the 88 depleted shRNAs. **D)** Stacked bar graph showing the frequency of each nucleotide in the 7mer seed for the 88 depleted and for all other shRNAs (Top). P < 0.05 was considered a significant association as calculated by Pearson’s chi-square test. List of 6mer seeds that appear multiple times within the 7mer seeds of the selected 88 depleted shRNAs (Bottom).

### Depleted shRNAs reduce viability and growth of prostate cancer cells and xenografts

To validate and more thoroughly determine the effect of the depleted shRNAs on PCa cell growth and viability, we used siRNAs with guide strand sequences identical to eight selected depleted shRNAs out of the initial 88 (siRNA mimics) and determined their effects on PCa cell growth and viability. Two non-target siRNAs (siNT-1 and siNT-2, see materials and methods for non-target seed sequence selection) and “no siRNA” were used as negative controls. LNCaP and 22Rv1 cells transfected with any of the siRNA mimics showed significantly reduced growth (Figures 2A-B) and viability (Figures 2E-F) to varying degrees, with respect to cells not transfected, or transfected with the control siRNAs, and induced PARP cleavage and H2AX phosphorylation, markers of cell death and DNA damage, respectively (Figures 2I-J). Since the AR is essential for PCa cell growth, we also analyzed the effect of these siRNAs on the expression of the AR and a representative AR target. Several siRNA mimics reduced the expression of the full-length AR, AR-V7, and androgen responsive proteins, in a cell line specific manner (Figures 2I-J). However, transfection of the same cell lines with an siRNA pool that targets the AR, and strongly reduced its expression (Figures 2I, 2J), was significantly less effective at reducing growth and viability compared to the siRNA mimics (Figures 2A-B, 2E-F), indicating that the effect of our select siRNAs is not just a consequence of their ability to target the AR. In fact, some of the selected siRNAs (e.g. siUACUGGC) did not affect AR expression, but still significantly reduced expression of PSA (an androgen responsive protein), as well as growth and viability of LNCaP cells (Figures 2A, 2E, 2I).

**Figure 2:**
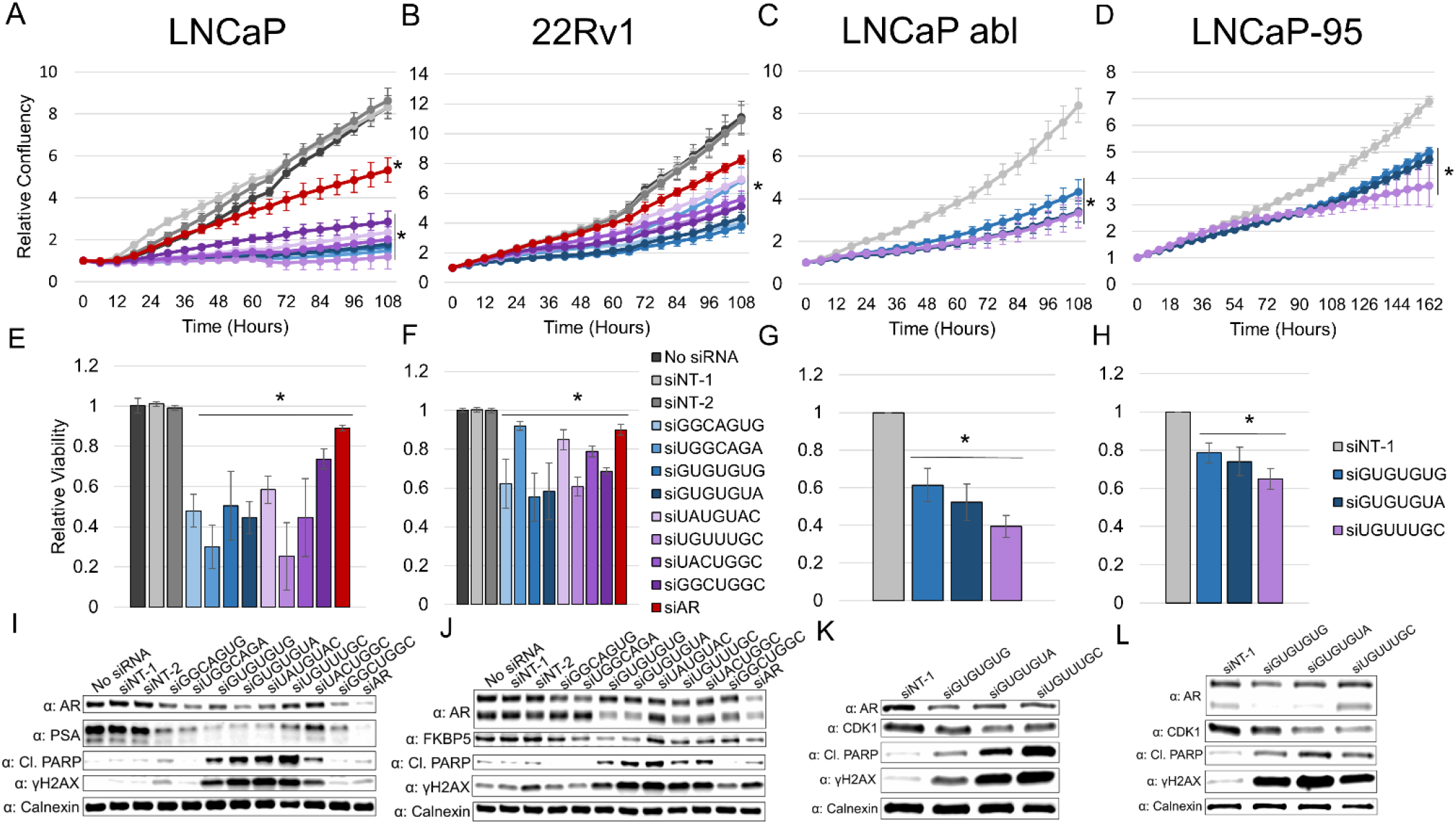
Validation of toxic RNAi sequences. Cell growth (A-D) and viability (E-H) graphs, and western blots (I-L) from lysates, of LNCaP (A,E,I), 22Rv1 (B,F,J), LNCaP abl (C,G,K) and LNCaP-95 (D,H,L) cells transfected with the designated siRNAs. Relative confluency was used as a surrogate for cell growth and was determined via IncuCyte®. Relative viability was assessed by trypan blue 6 days after siRNA transfection (8 days after transfection for LNCaP-95 cells). Western blots show AR (top band: AR full length, bottom band: AR-V7 in 22Rv1 and LNCaP-95), androgen responsive proteins (PSA, FKBP5, CDK1) and cell death and DNA damage markers (cleaved PARP and γH2AX,); lysates were collected 72 hours after siRNA transfection and Calnexin was used as a loading control (I-L). N = 3 or 4 for each experiment, * q < .05 compared to pooled siNT-1, siNT-2 and no-siRNA (A, B, E, F) or siNT-1 (C, D, G, H) transfection was considered significant as determined by t-test.

Three siRNAs that consistently demonstrated potent effects on growth and viability of LNCaP and 22Rv1 cells (siGUGUGUG, siGUGUGUA, siUGUUUGC) were further validated in additional PCa cell lines. Transfection with any of these three siRNAs decreased growth (Figures 2C-D) and viability (Figures 2G-H) of LNCaP abl and LNCaP-95 cells to varying degrees, and induced PARP cleavage and H2AX phosphorylation (Figures 2K-L). They also reduced expression of AR and AR-targets. siGUGUGUG and siGUGUGUA were shown to reduce the expression of AR-V7 in LNCaP-95 cells (Figure 2L), similar to the results seen in the AR-V7+ 22Rv1 cells. The same three siRNAs also reduced the growth of 22Rv1 cells cultured in the presence or absence of DHT to a similar degree (Figure S4). These data suggest that the identified siRNAs are toxic across different PCa cell lines and growth conditions.

We then examined the effect of one of the toxic shRNAs on the growth of subcutaneous tumors in mice. To this end, we cloned the shRNA containing the UGUUUGC seed sequence or a shNT control in the Dox inducible lentiviral vector used for library construction and expressed them in 22Rv1 cells. In cell culture, Dox treatment significantly reduced growth and viability of 22Rv1 cells expressing the shUGUUUGC shRNA compared to shNT expressing cells (Figures S5A-B). For in vivo experiments, these two cell lines were grown in the absence of Dox –to prevent cell death, mixed with 50% basement membrane extract and inoculated subcutaneously into flanks of NRG mice that were pre-fed (2 days) and kept in a Dox containing-diet for the duration of the experiment. Consistent with the *in vitro* observations, expression of the shUGUUUGC shRNA, significantly reduced tumor growth in this murine model (Figure S5C). These results validate our novel seed-based shRNA screen as a method to identify toxic RNAi molecules that inhibit growth and viability of PCa cells. Importantly, while many of these sh/siRNAs reduce AR expression, they do not exclusively rely on, and result in more potent viability/growth effects than, targeting the AR alone.

### Depleted shRNAs seed sequences are predicted to target AR coregulatory and prostate cancer essential genes

The selected toxic sh/siRNAs only differ among them, and from non-toxic ones, in the 7mer seed sequence. Therefore, we anticipated that the toxic effect of the selected sh/siRNAs was mediated by the seed sequence, as we and others have previously shown for other small toxic RNAs.^17–19^ To confirm this prediction, we designed siRNAs derived from the toxic siGUGUGUG, siGUGUGUA or siUGUUUGC, in which the seed sequence was shifted out of the seed region, or the order of nucleotides within the seed region was altered (Figure S6C). Transfection of LNCaP cells with these modified siRNAs did not affect their growth or viability, as compared to the non-target siRNA control (siNT-1; Figure S6A, S6B), suggesting that altering the seed reverses the toxic phenotype, and therefore that the observed toxic phenotypes are seed mediated. These results supported the use of the miRNA database, miRDB (www.mirdb.org) (which uses the seed sequence) to identify the predicted targets of the selected 88 depleted shRNAs.

Based on our previous work^17^ and the essential role of the AR in PCa cell growth, we hypothesized that AR coregulatory and PCa essential genes would be enriched among the predicted targets of the depleted seeds. In fact, the predicted targets of 32 out of the 88 toxic shRNAs were significantly enriched for AR coregulatory^25^ and LNCaP essential^26^ genes (Figure 3A, Table S1). More broadly, analysis with the predicted targets of all the 88 toxic shRNAs indicated a more significant enrichment of AR coregulatory^25^ and LNCaP essential genes^26^ than of the non-PCa essential^18,27,28^ genes (Figure S7A) and the significance of their enrichment (log10q values) were strongly correlated (Figure S7B). The enrichment of AR coregulatory and LNCaP essential genes (log10q) within the predicted targets of the 88 toxic shRNAs was also correlated with their respective target scores (Figure S8A-B). The Target Score represents the abundance of sequences complementary to the seed sequences of the toxic shRNAs in the 3’UTRs of AR coregulatory or LNCaP essential genes, relative to their abundance in the 3’UTRs of the whole transcriptome. More strikingly, a positive correlation was found between the AR coregulatory and the LNCaP essential genes target scores when considering the sequences complementary to the seeds of all the shRNAs in the screen (n= 15,572; Figure S8C). This suggests that the similar abundance of specific 6 and 7 nucleotide sequences in the 3’UTR may account for the paired enrichment of AR coregulatory and PCa essential genes observed within the predicted targets of the depleted shRNAs. Importantly, for the depleted shRNAs, predicted targeting of the AR coregulatory genes is significantly associated with predicted targeting of the *AR* (Figure 3B), suggesting that RNAi of hormone receptors and their coregulators may happen concurrently.

**Figure 3:**
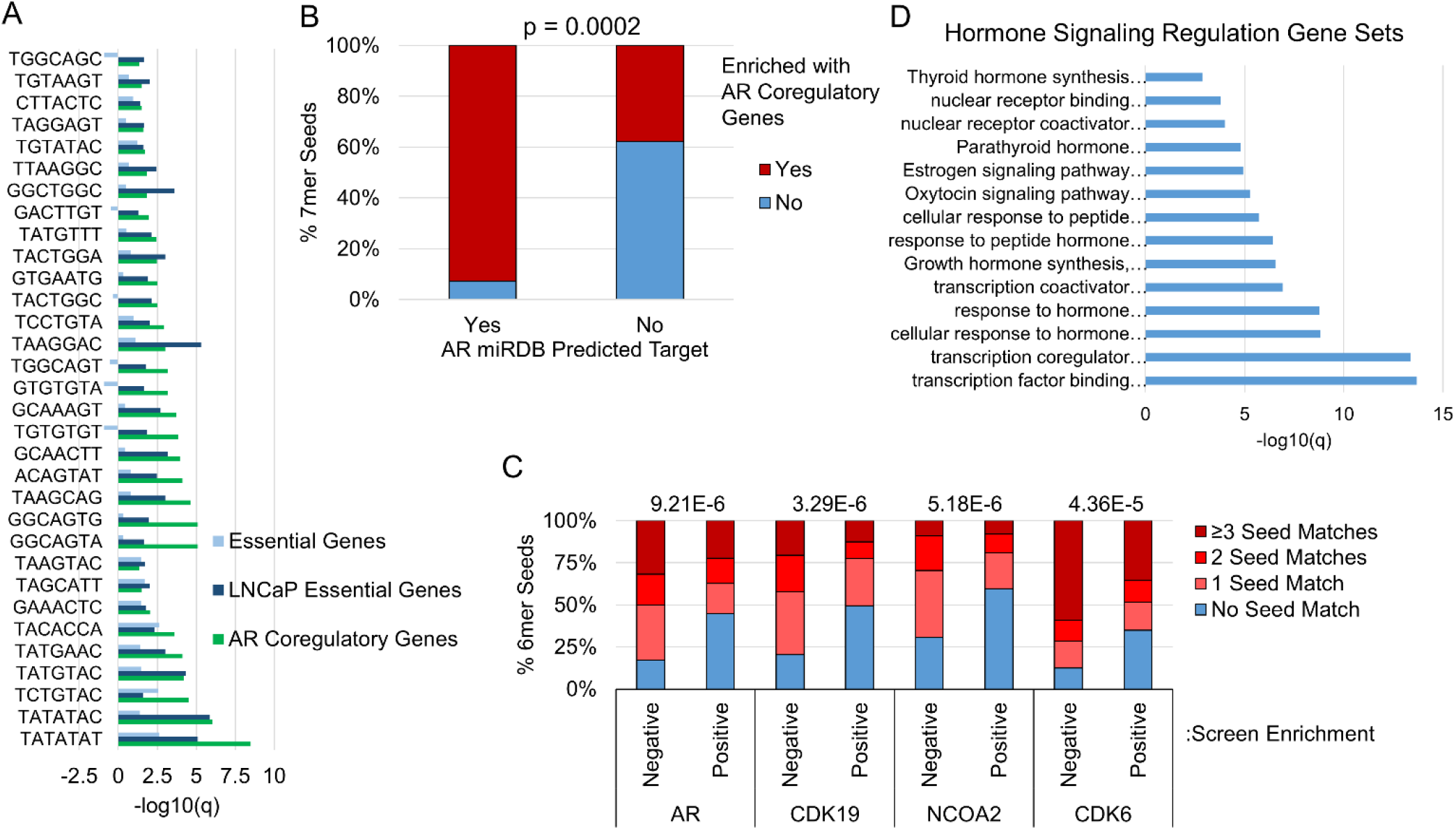
Depleted shRNAs are predicted to target AR coregulatory and prostate cancer essential genes. A) Bar graph showing the -log10(q) of enrichment of AR coregulatory, LNCaP essential (PCa essential genes) and essential (obtained from non-PCa cells) genes among the predicted targets for depleted shRNAs. Only the ones with significant AR coregulatory and LNCaP essential gene enrichment are shown (n= 32). shRNAs are named by their 7mer seed sequence. Negative values indicate that gene sets are under-represented among predicted targets. –log10(q) > 1.3 (q < 0.05) was considered significant, as determined by hypergeometric distribution. B) Stacked bar graph showing that a high percentage of depleted shRNAs predicted to target the AR, are also predicted to target AR coregulatory genes (red). P < 0.05 was considered a significant association as calculated by Fisher’s exact test. C) Stacked bar graph showing the percentage of negatively selected (depleted, n = 88) and positively selected (enriched, n = 474) shRNAs with 0, 1, 2 or 3 or more 6mer seed matches (complementarity) in the 3’UTR of each of the indicated target genes (AR, CDK19, NCOA2, CDK6). P-values are above the plot for each association, and p < 0.05 was considered a significant association as calculated by Pearson’s chi square test. D) Bar graph showing the significance (–log10q) of GO and reactome gene sets involved in hormone signaling regulation enriched among genes predicted to be targeted by > 10% of depleted seeds. GO and reactome gene sets were analyzed using www.metascape.org. –log10(q) > 1.3 (q < 0.05) was considered significant, as determined by hypergeometric distribution.

Genes predicted to be targeted by at least 10% (≥ 9) of the depleted shRNAs, were also enriched in AR coregulatory and LNCaP essential genes (Figure S9), and included several genes known to be important for PCa cell growth and survival, for example the *AR*, the AR coregulators *NCOA2* and *CDK6,*and the LNCaP essential gene *CDK19.* Seed sequence complementarity in the 3’UTR of these 4 genes was more strongly associated with depleted shRNAs than with positively selected shRNAs (Figure 3C). Moreover, genes targeted by 9 or more depleted shRNAs were also enriched in several hormone signaling GO and reactome gene sets (Figure 3D), suggesting that hormone receptor signaling inhibition is a common mechanism through which RNAi seed sequences reduce growth and/or viability of hormone driven cancer cells.

### Downregulation of AR coregulatory and prostate cancer essential genes is associated with sequences in their 3’UTR complementary to the GUGUGUA and UGUUUGC seeds

To confirm downregulation of AR coregulators and PCa essential genes by the toxic siRNAs, we conducted RNA seq with RNA from 22Rv1, LNCaP and LNCaP abl cells, transfected with siNT-1, and the toxic siGUGUGUA or siUGUUUGC siRNAs, and analyzed gene expression changes. RNA was also obtained from LNCaP abl and 22Rv1 cells after transfection with AR-targeted siRNA (siAR), but not for LNCaP, since AR regulated genes in LNCaP cells are well defined in the literature and reported by us.^17^RNA was extracted prior to loss in viability triggered by the toxic siRNAs, which was at 40 hours post-transfection in LNCaP cells, 48 hours for LNCaP abl and 64 hours for 22Rv1 cells (data not shown). Genes that exhibited ± 0.5 log2 fold change and adjusted p-value <0.05 in samples from cells transfected with toxic siRNAs relative to siNT control were considered to be significantly differentially expressed genes (DEGs).

Consistent with the data from the predicted target analyses using the miRDB software (Figure 3A), the RNA seq data showed that genes downregulated by siGUGUGUA and siUGUUUGC in LNCaP and other PCa cell lines were significantly enriched in AR coregulatory and LNCaP essential genes (Figure S10). To verify that downregulation occurs via a seed mediated mechanism, we performed a series of tests: i) we used the cWords software^29^ to identify 6-7 nucleotide 3’UTR sequences that were most associated with mRNA downregulation in the PCa cells transfected with siGUGUGA, siUGUUUGC, and then determined whether the enriched sequences were complementary to the seed sequence of those siRNAs. The analyses demonstrated that the 6-7 nucleotide 3’UTR sequences most significantly associated with downregulated genes were in fact complementary to the siGUGUGUA, siUGUUUGC seeds, ii) The Kolmogorov–Smirnov (KS) test was used to confirm that genes with 3’UTR 7mer complementary sequences to the GUGUGUA or UGUUUGC seeds were significantly downregulated by the respective siRNAs, and iii) GSEA showed that miRDB predicted targets (which are based on the 7mer seed sequence) for both siRNAs were significantly downregulated (Figures 4A-D, S10A-D, S11A-D, S12A-D, S13A-D). Similarly, seed sequence complementarity in the 3’UTR was significantly associated with AR coregulatory and LNCaP essential gene downregulation (Figures 4E, S11E, S12E, S13E). These data show that siGUGUGUA and siUGUUUGC downregulate multiple AR coregulatory and PCa essential genes through a seed mediated mechanism.

**Figure 4:**
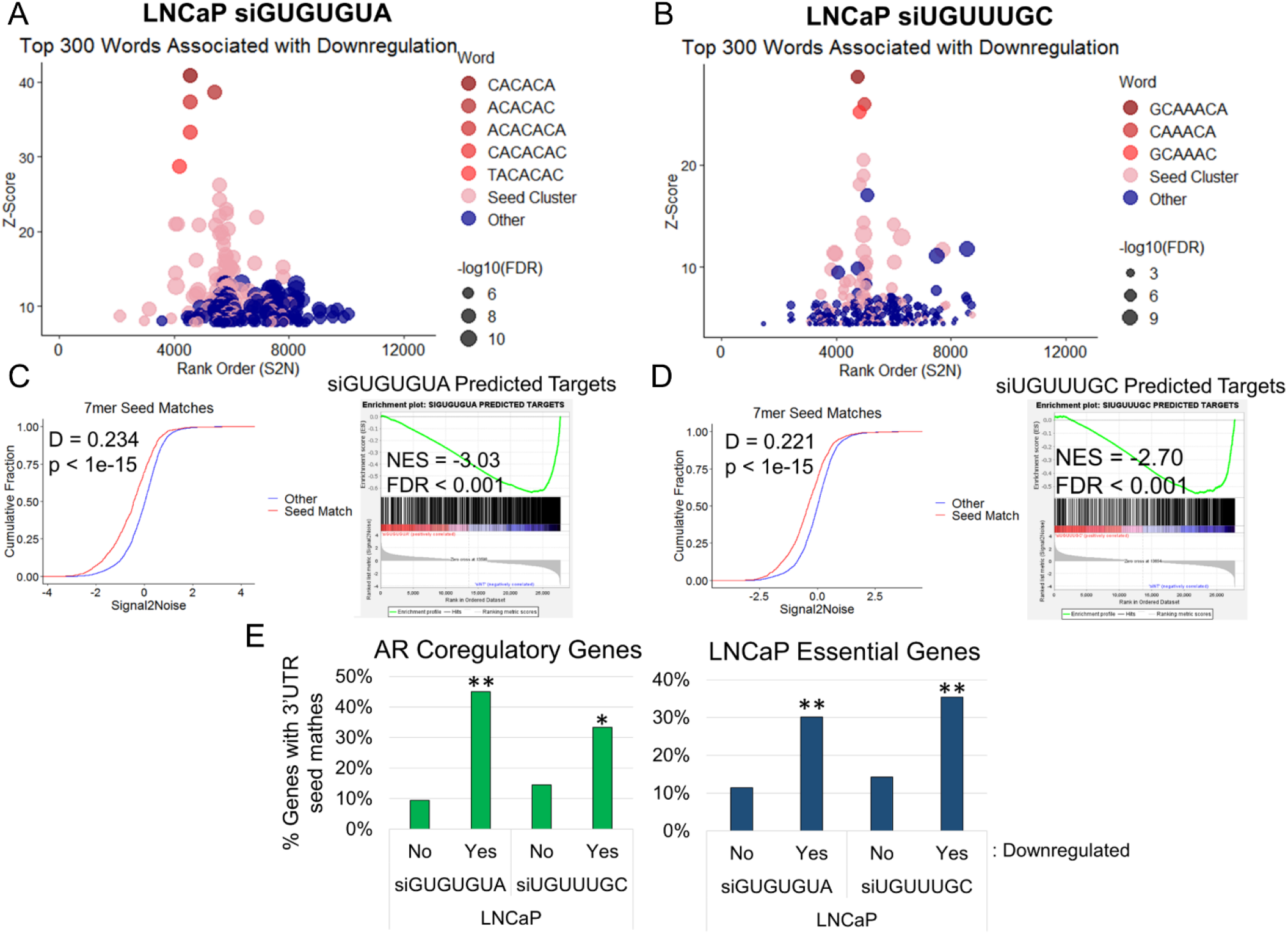
Global, AR coregulatory and prostate cancer essential gene downregulation in LNCaP cells is associated with sequences complementary to the GUGUGUA and UGUUUGC seeds in the 3’UTR. (A-B) cWords plots displaying the top 300 words (6 and 7 nucleotide 3’UTR sequences) associated with gene downregulation in LNCaP cells transfected with siGUGUGUA (A) or siUGUUUGC (B). The top seed-related sequences are indicated; “seed cluster” includes other seed-related sequences. Genes were ranked ordered by signal to noise values from GSEA. (C-D) eCDF plots from LNCaP cells transfected with siGUGUGUA (C, left) or siUGUUUGC (D, left) showing the cumulative fraction of genes across the signal to noise rank ordered gene expression list, for genes with and without 7mer seed match sequences in their 3’UTR; p < 0.05 as determined by KS test, D and p values are labeled on plots. GSEA enrichment plots for miRDB predicted target gene sets in LNCaP cells transfected with siGUGUGUA (C, right) or siUGUUUGC (D, right). E) Bar graphs showing the percentage of AR coregulatory (Left) and LNCaP essential (Right) genes with 7mer and/or multiple 6mer seed matches to siGUGUGUA or siUGUUUGC in the 3’ UTR, stratified by whether the genes are downregulated (Yes) or not (No) by the corresponding siRNAs in LNCaP cells. *p < 0.05, **p < 0.0001, as determined by fisher’s exact test.

### siGUGUGUA and siUGUUUGC siRNAs reduce the expression of androgen responsive and cell cycle genes in prostate cancer cells

Because AR coregulatory and essential gene networks were direct targets of siGUGUGUA and siUGUUUGC, we hypothesized that siGUGUGUA, siUGUUUGC and siAR would elicit similar changes in gene expression. Indeed, the RNA-Seq data indicated that a large number of upregulated and downregulated DEGs were common in response to siGUGUGUA, siUGUUUGC and siAR transfection in all cell lines (Figures 5A-B, S14A-B). However, downregulated DEGs generally demonstrated a stronger overlap than upregulated genes, suggesting that siGUGUGUA and siUGUUUGC may block the activation of similar genes and pathways, including AR-mediated gene activation. We next used GSEA to identify pathways affected by the toxic siRNAs. GSEA using the HALLMARK gene sets within the molecular signature database (http://www.gsea-msigdb.org) indicate that siGUGUGUA and siUGUUUGC significantly downregulate androgen signaling (HALLMARK_ANDROGEN_RESPONSE) and several cell cycle related gene sets (HALLMARK_E2F_TARGETS, HALLMARK_G2M_CHECKPOINT and HALLMARK_MITOTIC_SPINDLE) (Figures 5C-D, S14C-D). Furthermore, GSEA with other AR-signaling related gene sets, such as, AR Score^30,31^, AR-V7 regulated genes^32^, genes upregulated and downregulated by siAR in 22Rv1 and LNCaP abl cells in our current RNA seq, and genes upregulated and downregulated by DHT in LNCaP control cells from a previous publication by our lab^17^, showed significantly altered expression in siGUGUGUA and siUGUUUGC transfected cells in all cell lines (Figures 5E-F, S14E-F, S15A-B, S16A-B, S17A-D).

**Figure 5:**
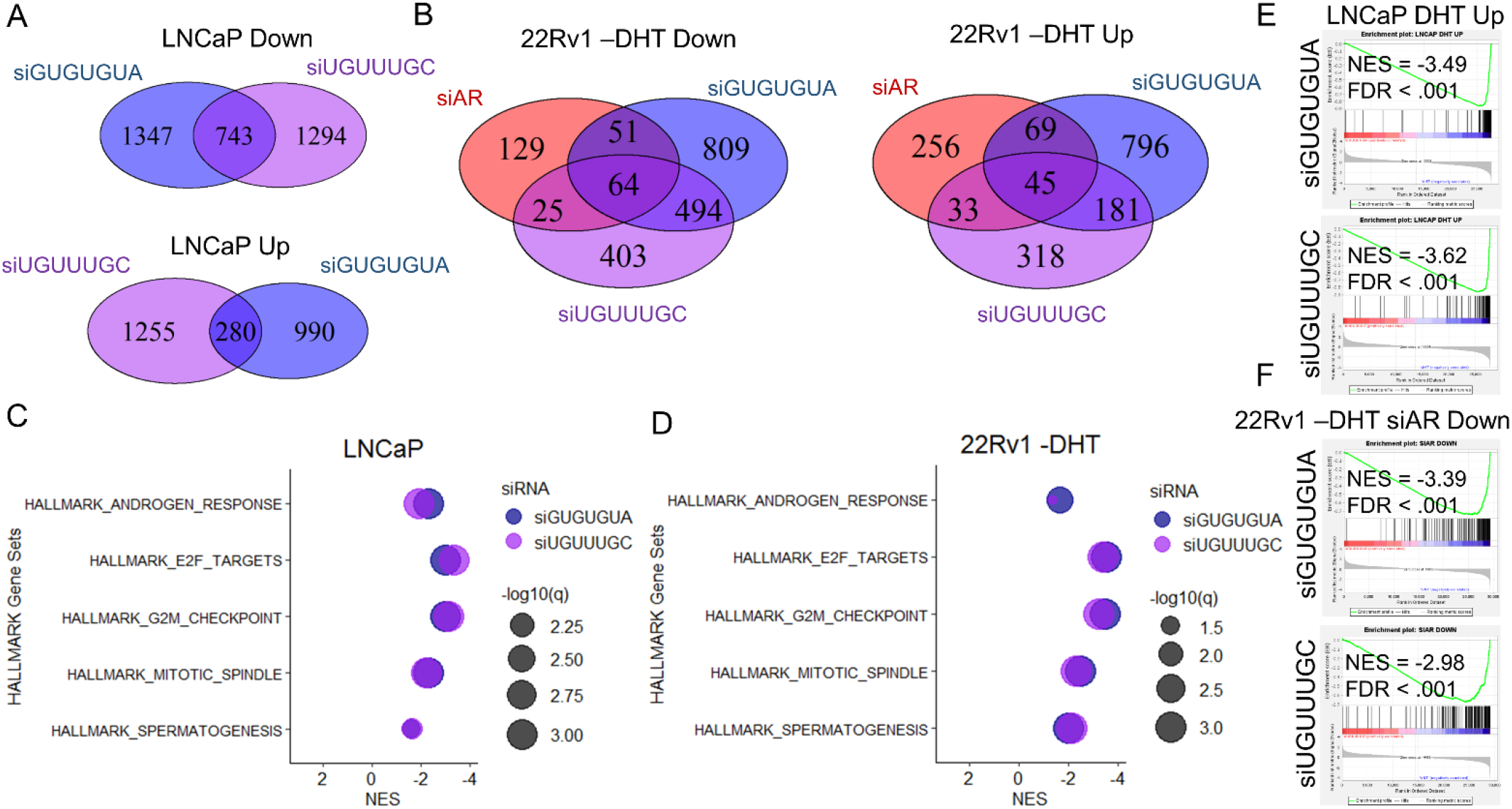
siGUGUGUA and siUGUUUGC reduce the expression of androgen responsive and cell cycle genes in PCa cells. (A-B) Venn diagrams showing the overlap of significantly upregulated and downregulated genes as determined by RNA-seq ([log2(FC)] > 0.5, padj < 0.05) in LNCaP cells (A) and 22Rv1 cells grown in the absence of DHT (B) transfected with siGUGUGUA, siUGUUUGC or AR targeted siRNA (siAR), compared to cells transfected with siNT-1. (C-D) Plots showing NES and –log10(q) for significantly upregulated and downregulated Hallmark gene sets in response to transfection with siGUGUGUA and siUGUUUGC in LNCaP (C) and 22Rv1 (D) cells. (E-F) GSEA enrichment plots showing the enrichment of the “LNCaP DHT up” gene set in LNCaP cells (E), and the enrichment of the “22Rv1 -DHT siAR down” gene set in 22Rv1 - DHT cells (F) transfected with siGUGUGUA and siUGUUUGC. Positive and negative NES indicate upregulated and downregulated gene sets respectively. FDR < 0.1 was considered significant.

Using RT-qPCR, we validated the significant downregulation of several AR responsive and cell cycle genes, as well as AR and AR coregulatory genes, in LNCaP, LNCaP abl and 22Rv1 cells transfected with the siGUGUGUA and siUGUUUGC siRNAs. (Figures S18A-B, S19A-B). Importantly, the AR and AR coregulatory genes contained 3’UTR sequences complementary to the seeds, while the AR regulated and cell cycle genes did not (Figures S18C and S19C). Together these data suggest that siGUGUGUA and siUGUUUGC directly target androgen signaling regulatory networks to indirectly inhibit androgen signaling modulate and cell cycle in ADPC and CRPC cells.

### Toxic siRNAs reduce prostate cancer cell growth and viability more effectively than the knockdown of AR or individual AR coregulatory genes and inhibit pre-established prostate cancer xenografts

Our data show that siGUGUGUA and siUGUUUGC target networks of AR coregulatory and essential genes. We therefore, hypothesized that siGUGUGUA and siUGUUUGC would inhibit the growth and viability of prostate cancer cells more significantly than siRNAs that target individual AR coregulators. To test this hypothesis, LNCaP and 22Rv1 cells were transfected with siRNAs targeting the AR coregulators p300 (*EP300*), HOXB13, and MLL (*KMT2A*), with the toxic siGUGUGUA, siUGUUUGC or siNT-1. siRNA targeting the AR was used as a control since we previously showed that is less efficient than the siGUGUGUA or siUGUUUGC siRNAs at inhibiting growth and viability of PCa cells (Figures 2A-B, 2E-F). Both siGUGUGUA and siUGUUUGC reduced the growth and viability of LNCaP and 22Rv1 cells more efficiently than the individual targeting of AR or AR coregulators (Figures 6A-D). RTq-PCR analysis demonstrated that siGUGUGUA and siUGUUUGC were in general more effective at reducing the expression of androgen responsive genes in LNCaP cells than the targeting of individual coregulators (Figures 6E). Interestingly, siGUGUGUA and siUGUUUGC also significantly decreased the expression of *EP300, HOXB13* and *AR* at different levels (Figures 6F).

**Figure 6:**
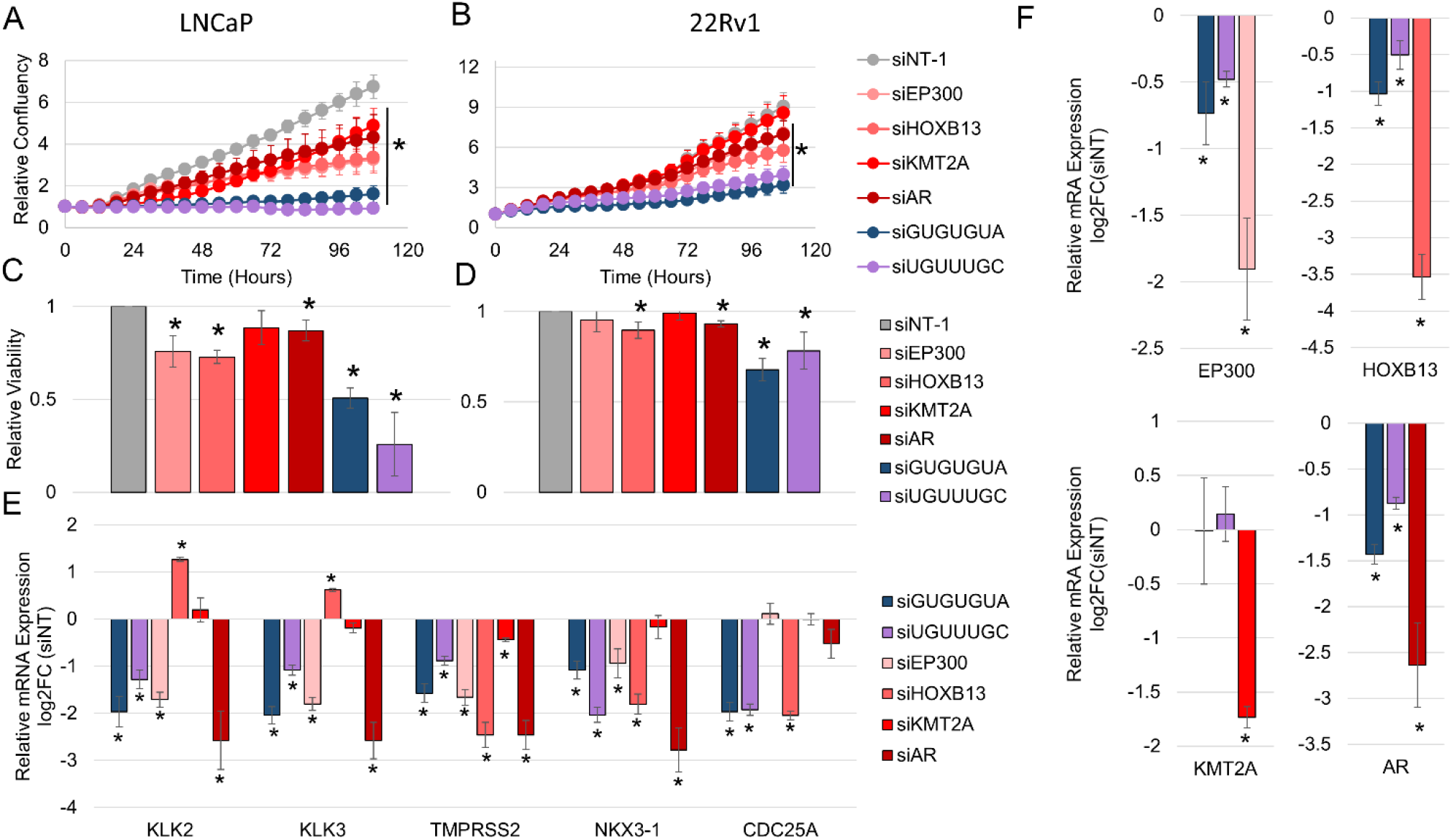
siGUGUGUA and siUGUUUGC reduce prostate cancer cell growth and viability more effectively than the knockdown individual AR coregulatory genes. (A-D) Cell growth (A-B) and viability (C-D) of LNCaP (A,C) and 22Rv1 (B,D) cells transfected with the indicated siRNA. Relative confluency was used as a surrogate for cell growth and was determined via IncuCyte. Relative viability was assessed by trypan blue 6 days after siRNA transfection. (E-F) Relative mRNA expression of androgen responsive genes, AR coregulators and AR in response to transfection of the indicated siRNA in LNCaP cells. Relative viability and gene expression are expressed as fold and log2(fold) of siNT-1 respectively. *q < 0.05 was considered significant as determined by t-test, n=3.

Importantly, while transfection with siGUGUGUA and siUGUUUGC negatively affected PCa cell growth and viability, they had negligible effects on the growth and viability of benign prostate cell lines, NhPre1 and BhPre1 (Figure S20). This suggests that the seed-mediated toxicity is specific to cancer cells, as previously described,^17,33^ which is a desirable therapeutic characteristic. In order to determine the therapeutic potential of this mechanism, we examined the effect of toxic shRNAs on the growth of pre-established tumors. To this end, we established xenografts with 22Rv1 cells expressing the Dox inducible shUGUUUGC or shNT as control. After xenografts reached a volume of approximately 120 mm^3^ mice were switched to a Dox containing diet. Dox induction of shUGUUUGC significantly reduced xenograft growth rate when compared to xenografts expressing shNT (Figure S21), suggesting that shUGUUUGC can inhibit pre-established growth of advanced PCa *in vivo*. Overall our data suggest that through a multi-target mechanism, siRNAs like siGUGUGUA and siUGUUUGC may offer a therapeutic advantage over the targeting AR or individual AR coregulators in the treatment of PCa.

## DISCUSSION

Development of resistance to AR signaling targeted therapies is the main cause of PCa related mortality and the reason PCa remains the second leading cause of cancer related death in men.^1,2^ Identification of novel therapeutic modalities that prevent and/or cure resistant disease are therefore essential for the effective management of PCa. The broad range of adaptations that account for development of resistance to AR-signaling targeted therapies, have prompted recent clinical trials that asses the efficacy of double or triple combination therapies for castration sensitive and CRPC.^34–39^ The results of most of these trials are currently limited, but some demonstrated a significant overall survival benefit.^34,35^ While these developments point to the promise of combination therapy for the treatment of this disease, they also underline the difficulties of finding the appropriate combinations for maximum efficacy and limited interactions.

AR coregulators modulate AR-transcriptional output and therefore simultaneously targeting multiple coregulators, alone or in combination with AR directed therapies, could represent a very effective synergistic therapeutic approach for PCa. Unfortunately, the functional redundancy among coregulators, our incomplete knowledge of the AR-coregulatory arsenal and function, and the lack of specific inhibitors complicate this approach. A potential path to overcome these limitations is the use of a seed-mediated RNAi approach. Similar to miRNAs, a single exogenous small RNA can target multitude of mRNAs via their seed sequence,^40–42^ ultimately preventing the expression of the corresponding proteins. While miRNAs often have a modest effect on individual targets, the effect is amplified due to the fact that the targets of a single miRNA frequently encode proteins that participate in common pathways,^15,16^ resulting in an additive or synergistic effect. This aspect is especially relevant for PCa where growth and survival of cancer cells is largely dominated by the AR-signaling pathway.

We began assessing the potential for seed-mediated RNAi as a therapeutic approach for PCa, by running a negative selection screen using a novel shRNA library expressing all the potential 7mer seed sequences, --within an otherwise identical stem-loop backbone, to identify those shRNAs with deleterious effect to PCa cells. Using this unbiased methodology, we were able to identify 88 shRNAs with distinct 7mer seed sequences whose expression in PCa cells led to growth inhibition and/or loss of viability (depleted from the population). Predicted target analyses demonstrated an enrichment for AR coregulatory and hormone signaling related gene sets among the genes predicted to be targeted by numerous depleted shRNAs, supporting the hypothesis that androgen signaling regulatory networks can be targeted by RNAi to inhibit the growth of PCa cells. Furthermore, siRNA mimics with two selected seed sequences, GUGUGUA and UGUUUGC, were shown to: 1) reduce PCa cell growth and viability *in vitro*, 2) lower the expression of the AR (including AR-V7), as well as AR coregulatory and essential gene networks associated with 3’UTR sequences complementary to the seed and 3) inhibit global androgen signaling and cell cycle gene expression. *In vivo*, expression of a shRNA with the UGUUUGC seed sequence, reduced xenograft tumor growth. The effects in PCa cells were evident in both ADPC and CRPC cells. Importantly, we demonstrated that both siGUGUGUA and siUGUUUGC reduced PCa cell growth and viability to a greater extent than targeting AR or individual AR coregulators using siRNA. These data support the validity and potential for using RNAi to target networks of AR coregulatory and essential genes for the treatment of PCa.

Several studies have investigated AR targeted microRNAs (miRNAs) in PCa,^24,43,44^ but little work has been done exploring the potential of miRNAs to modulate androgen signaling regulatory networks. One study revealed that several tumor suppressive miRNAs, that are downregulated in CRPC, directly target the AR in addition to some of the NCOA family coregulators.^45^ Given the function of miRNAs to target numerous genes, we hypothesize that some endogenous miRNAs may function to regulate hormone signaling through lowering the expression of receptors and coregulators. In fact, several of the siRNAs with seed sequences identified in our screen and that we validated to reduce PCa cell growth and viability (Figure 2) share seed sequences with known miRNAs: miR-34, GGCAGUG; miR-6867-5p, GUGUGU; miR-7150, UGGCAG; miR-3924, UAUGUA; miR-3652/4430/4505/5787, GGCAGG. In addition to targeting the AR, siRNAs containing all of the above seeds showed an enrichment of AR coregulatory genes among their predicted targets. In fact, while, miR-34 has been shown to have a role in PCa through regulating AR expression,^23,24^ our data suggest that it regulates androgen signaling likely through targeting several AR coregulators as well. Together our results support the concept that miRNAs control hormone signaling networks.

The success of our approach relies on the ability of small RNAs to directly affect the expression of many target mRNAs by acting essentially as microRNAs (miRNAs). While the main caveat of this kind of approach is the potential for unwanted toxicities, i.e. immune stimulation^46–49^, our results indicated that the selected siRNAs have minimal effects on the growth/viability of benign prostate cells, suggesting minimal toxicity and confirming previous results.^17,33^ We did not observe toxicity in studies in which the seed sequence was moved out of place or rearranged (Figure S6) while still conserving putative immunostimulatory motifs.^46–48^ While this suggests that the selected siRNAs do not promote immune stimulation in prostate cells, strategies to mitigate this effect have been described and could be implemented.^50–53^ Moreover, most toxic shRNAs identified in our screen do not seem to primarily target general essential genes, as we didn’t observe enrichment of the non-PCa essential gene set within their predicted targets. In fact, when considering the group of 8 siRNAs selected to validate our screen, there was no correlation between their effect on the observed PCa cell viability and on the viability of other cancer cells as determined in silico using the 6merdb (www.6merdb.org).^19^ This points to PCa specific toxicity for a subset of the selected siRNAs.

While our approach was aimed to identify therapeutics for PCa, its applications go far beyond. It is possible that many of the sequences identified in the screen could also have therapeutic potential in other hormone driven cancers, such as breast and ovarian cancers. In fact, several hormone signaling regulatory gene sets involving different receptors (i.e. thyroid, parathyroid, oxytocin and estrogen receptors), were enriched among genes predicted to be targeted by several depleted shRNAs (Figure 3D), likely due to similarity in RNAi relevant sequences in the 3’UTR of hormone signaling regulatory genes. Similarly, this novel screen could be conducted in cell lines of other cancer types, in order to identify potentially therapeutic seed sequences. Finally, since our screen revealed enrichment of androgen signaling regulatory networks, it is possible that predicted target analyses from the depleted shRNAs in combination with other biochemical and bioinformatic approaches could be helpful to identify novel AR coregulators.

In summary, in this study, using a novel seed-based shRNA screen in PCa cancer cell lines, we identified small RNAs that inhibit androgen signaling through seed-mediated targeting of androgen regulatory and essential gene networks, resulting in a reduction in cell growth and viability. These results warrant further investigation into the use of RNAi to target hormone signaling networks as a potential therapeutic strategy for PCa.

## MATERIALS AND METHODS

### Cell culture

LNCaP (CRL-1740™) and 22Rv1 (CRL-2505™) cell lines were obtained from American Type Culture Collection (ATCC®, Manassas, VA, USA). LNCaP abl cells were obtained from Dr. Zoran Culig (University of Innsbruck). LNCaP-95 cells were obtained from Dr. Jun Luo (from the Genetic Resources Core Facility at Johns Hopkins University). BHPre1 and NHPre1 cells were obtained from Dr. S. Hayward (NorthShore Research Institute). LentiX-293T packaging cells were obtained from Clontech/Takara Bio (Mountain View, CA). None of these cell lines are on the list of contaminated and misidentified cell lines reported by ICLAC (https://iclac.org/databases/cross-contaminations/). Cells tested negative for mycoplasm using the Mycosensor PCR assay kit (Agilent, Santa Clara, CA, USA) or the LookOut Mycoplasma PCR detection kit (Sigma-Aldrich, St. Louis, MO, USA). LNCaP abl were subjected to short tandem repeat (STR) DNA profiling by IDEXX BioAnalytics (Columbia, MO).

LNCaP abl and LNCaP-95 cell lines were maintained in RPMI 1640 phenol red free medium (Gibco, Gaithersburg, MD, USA) supplemented with 10% charcoal-stripped serum (CSS) (Corning, Corning, NY, USA), 100 units/mL penicillin, 100 μg/mL streptomycin, Amphotericin B, and 2 mM L-glutamine. LNCaP and 22Rv1 cell lines were maintained in RPMI Glutamax growth media (Gibco) supplemented with 10% fetal bovine serum (FBS) (Corning, Corning, NY, USA), 100 units/mL penicillin, 100 μg/mL streptomycin, Amphotericin B, and 2 mM L-glutamine. BHPre1 and NHPre1 cell lines were maintained in HPrE-conditional medium, as previously described.^54^ For growth in presence or absence of dihydrotestosterone (DHT; 1 nM or 10 nM DHT; 0.0001% or 0.00001% EtOH vehicle control) 22Rv1 cells were maintained in RPMI Glutamax growth media supplemented with 10% CSS.

### shRNA library

The custom lentiviral shRNA library (Cellecta, Mountain View, CA, USA) consisted of 15,572 unique shRNAs. It contains shRNAs with all possible combinations of 7mer seed sequences (except for 812 7mer sequences that could not be cloned due to restriction enzyme and Pol III transcription termination sequence incompatibility). The rest of the sequence is common for all shRNAs (See Figure 1A for shRNA sequence), and the stem loop has previously been shown to yield consistent processing via Dicer;^55^ furthermore, the non-seed mature guide strand sequences have been previously demonstrated to be non-toxic.^19^ shRNAs were cloned in the pRSIT16-U6Tet-sh-CMV-TetRep-2A-TagRFP-2A-Puro vector (SVSHU6T16-L, Cellecta), that allows doxycycline induction. Each individual shRNA was linked to a unique sequence barcode for shRNA read identification with next generation sequencing (NGS).

### shRNA negative selection screen

For the shRNA screen, all cell lines were maintained in complete RPMI growth media, except for 22Rv1 cells which were grown in the presence and absence of 10 nM DHT for six days prior to transductions and throughout the screen. Cells were transduced with the shRNA library at a MOI of 0.35 and 500 fold representation (two repeats per cell line). After puromycin selection, half of the surviving cells (> 1,000 fold representation) were collected as “day 0” control samples, and stored at -80°C for genomic isolation at a later time. Flow cytometry was used to confirm that greater than 90 percent of cells were RFP positive. The rest of the cells were kept in low dose puromycin and 500 ng/ml doxycycline was added to induce shRNA expression. These cells were continuously passaged at 50-60% confluency to maintain a minimum of 1,000 fold representation for 10 doublings and then collected as the end point sample (for 22Rv1 –DHT and 22Rv1 +DHT cells we also obtained a sample after two doublings). Genomic DNA was extracted from at least 2×10^7^ cells (> 1,000 fold representation) from the “day 0” and the end point samples using DNeasy Blood and Tissue Kit (QIAGEN, Hilden, Germany) followed by DNA precipitation to concentrate the DNA. DNA samples were sequenced by NGS, and sequence reads were normalized as previously described.^21^ Z-scores were then calculated for shRNA normalized read counts in two and ten doubling samples relative to baseline (“day 0”). Significantly depleted shRNAs were then identified as shown in Figure S1. Average z-score > 0.75 and q < 0.3 across all cell lines was used as a cutoff for selecting a pool of positively selected shRNAs.

### Predicted target and 3’UTR sequence analyses

The miRNA database (www.mirdb.org) custom prediction tool was used for predicted target analyses. Enrichment of AR coregulatory^25^ LNCaP essential^26^ and essential^18,27,28^ gene sets among predicted targets was determined by hypergeometric distribution, and q-values were calculated to correct for multiple comparisons. Enrichment of GO and reactome gene sets was analyzed using metascape (www.metascape.org). Fasta files containing Ensembl 103 MANE select transcript 3’UTR sequences were obtained from ensembl Biomart, and seankross/warppipe (https://rdrr.io/github/seankross/warppipe/) and stringr R packages were used for sequence analyses. ARCTS, LEGTS and EGTS indicated the enrichment of 6mer and 7mer seed complementary sequences (seed match) in the 3’UTR of AR coregulatory,^25^LNCaP essential^26^ and essential^18,27,28^ genes, respectively, relative to their occurrence across the whole transcriptome (seed match/kb_expected_) (Z-Scores calculated from normalized residuals: (seed match/kbobserved-seed match/kb_expected_)/sqrt(seed match/kb_expected_)).

### siRNA transfections and lentiviral shRNA vector transductions

Unmodified custom siRNAs and ON-TARGET-Plus pool siRNAs were obtained from Dharmacon (Horizon Discovery, Waterbeach, UK). Non-target siRNA seeds were selected due to the low occurrence of complementary sequences in 3’UTRs across the transcriptome (Ensembl 103 MANE select transcript 3’UTR sequences) and their lack of toxicity in the screen.

#### Unmodified siRNA guide strand sequences

<u>siNT-1</u>: UCGUACGAACAUGUAACCG

<u>siNT-2</u>: UUUCGCGAACAUGUAACCG

<u>siGGCAGUG</u>: UGGCAGUGACAUGUAACCG

<u>siUGGCAGA</u>: UUGGCAGAACAUGUAACCG

<u>siGUGUGUG</u>: UGUGUGUGACAUGUAACCG

<u>siGUGUGUA</u>: UGUGUGUAACAUGUAACCG

<u>siUAUGUAC</u>: UUAUGUACACAUGUAACCG

<u>siUGUUUGC</u>: UUGUUUGCACAUGUAACCG

<u>siUACUGGC</u>: UUACUGGCACAUGUAACCG

<u>siGGCUGGC</u>: UGGCUGGCACAUGUAACCG

<u>siCGUGUUU</u>: UCGUGUUUACAUGUAACCG

<u>siNT-1-UGUGUGUG</u>: UCGUACGAUGUGUGUGACAΔ

<u>siCGCAUGU-GUGUG</u>: UCGCAUGUGUGUGACAUGU

ON-TARGET-plus pool siRNAs: *AR* (L-003400-00-0005), *EP300* (p300) (L-003486-00-0005), *HOXB13* (L-012226-00-0005), *KMT2A* (MLL) (L-009914-00-0005).

Dharmafect siRNA transfection protocol was used for transfections (https://horizondiscovery.com/-/media/Files/Horizon/resources/Protocols/basic-dharmafect-protocol.pdf). 30 nM siRNA was used for each transfection. 0.2% dharmafect reagent #3 was used for transfections in LNCaP, 22Rv1, LNCaP abl and LNCaP-95 cell lines. 0.2% dharmafect reagent #1 was used for transfections in BHPre1 and NHPre1 cell lines. For experiments involving DHT treatment, 22Rv1 cells were grown in the presence and absence of 1 nM DHT for six days prior to siRNA transfections.

Individual shRNAs were cloned into the pRSIT16-U6Tet-sh-CMV-TetRep-2A-TagRFP-2A-Puro vector (SVSHU6T16-L, Cellecta) using the following oligonucleotides:

shNT For:

ACCGGCGGTTACATGTTCGTACGACTCCTGACCCAAGTCGTACGAACATGTAACCGTTTTTGAA

shNT Rev: CGAATTCAAAAAACGGTTACATGTTCGTACGACTTGGGTCAGGAGTCGTACGAACATGTAACCGC shGUGUGUA For: ACCGGCGGTTACATGTTACACACACTCCTGACCCAAGTGTGTGTAACATGTAACCGTTTTTTGAA shGUGUGUA Rev:

CGAATTCAAAAAACGGTTACATGTTACACACACTTGGGTCAGGAGTGTGTGTAACATGTAACCGC shUGUUUGC For:

ACCGGCGGTTACATGTGCAAACAACTCCTGACCCAAGTTGTTTGCACATGTAACCGTTTTTTGAA shUGUUUGC Rev:

CGAATTCAAAAAACGGTTACATGTGCAAACAACTTGGGTCAGGAGTTGTTTGCACATGTAACCGC

Cells were transduced with lentiviruses containing the above plasmids at a MOI of 20, and were selected for seven days in 1 ug/ml puromycin.

### Cell growth and viability analyses

For cell growth assays, cells were seeded in 96 well plates at a concentration of 3,000 cells per well (LNCaP cells were seeded 4,000 cell per well) 24 hours after siRNA transfections or 48 hours after doxycycline treatment for cells expressing shRNAs. Cell growth was assessed using the IncuCyte Live Cell Imaging System. Percent confluence was measured every six hours, and values were normalized to percent confluence at the first reading (relative confluence). Trypan blue was utilized to calculate cell viability using the Nexcelom Cellometer Auto T4 after the completion of growth assays (132 or 186 hours after siRNA transfections, and 156 hours after doxycycline treatment for cells expressing shRNAs). Relative percent viability was calculated by dividing percent viability of cells transfected/transduced with experimental siRNA/shRNA by that of cells transfected/transduced with non-target control.

Statistics: two-tailed t tests were used to calculate significant differences in growth and viability. n= 3 or 4. p <0.05 was considered to be statistically significant (FDR (q) was calculated when indicated to correct for multiple comparisons, q < 0.05 was considered significant).

### Mouse xenografts

Animal procedures were approved by the Oklahoma Medical Research Foundation (OMRF) Institutional Animal Care and Use Committee (IACUC). Immunodeficient NRG (#007799) or NOD/Sci (#001303) mice (The Jackson Laboratory, Bar Harbor, ME, USA) were used for this study. 22Rv1 cells expressing dox-inducible shNT or shUGUUUGC shRNAs were mixed with 50% basement membrane extract (R&D Systems, Minneapolis, MN, USA) and injected subcutaneously into the flanks (3×10^6^ cell per injection). Mice were fed doxycycline-containing chow (200 mg/kg, Bioserv, Flemington, NJ) two days before the injection and for the remaining of the experiment and tumor volume was measured three times per week using calipers. Tumor volume was calculated using the formula ½ (Length × Width^2^). Significant differences in tumor growth between shUGUUUGC and shNT expressing cells were determined using a segmented linear mixed model. Specifically, we first identified the transition point and then we fitted a linear mixed model for the data before and after the transition point. R packages segmented, Ime4, ImerTest, nlme and emmeans were used. p < 0.05 was considered significant. For therapeutic experiments, after injections, mice were fed with normal chow until the tumors reach a volume of 100-120 mm^3^ and then transferred to Doxycycline containing diet until the end of the experiment.

### Western blot analysis and antibodies

Whole cell lysates were obtained using Cell Signaling lysis buffer (9803, Cell Signaling Technology, Danvers, MA, USA). Stain free mini-PROTEAN TGX gels (Biorad, Hercules, CA, USA) were used for SDS-PAGE protein separation, and proteins were transferred to 0.2 μm nitrocellulose membranes using the turbo transfer system (Biorad). Membranes were incubated in 5% nonfat dry milk (NFDM) in tris-buffered saline with 0.1% Tween 20 (TBST) for one hour for blocking, and then incubated overnight in primary antibodies diluted in 5% NFDM TBST. The following antibodies and dilutions were used: anti-PSA (76113, Abcam, Cambridge, UK) 1:1,000, anti-FKBP5 (2901, Abcam) 1:1,000, anti-CDK1 (A17, Abcam) 1:1,000, anti-AR (39781, Active Motif, Carlsbad, CA, USA) 1:1,000, anti-Cleaved PARP (Asp214) (5625S, Cell Signaling Technology) 1:1,000, anti-phospho-H2AX Ser139 (9718S, Cell Signaling Technology) 1:1,000, anti-Calnexin (22595, Abcam) 1:4,000.

Membranes were incubated for one hour in secondary antibody diluted in 5% NFDM TBST at a concentration of 80 ng/ml: Goat anti-rabbit or goat anti-mouse horseradish peroxidase (HRP)-conjugated secondary antibody (31460 and 31460 respectively, Thermo Fisher Scientific, Waltham, MA). Clarity Western ECL (Biorad) or SuperSignal West Femto (Thermo Fisher Scientific) and the ChemiDoc Touch Imaging System (Biorad) were used for chemiluminescent detection.

### qRT-PCR

RNeasy mini kit (QIAGEN) with on-column DNase incubation was used for RNA extractions. iScript reverse transcription supermix (Biorad) was used for reverse transcription reactions (500 ng RNA per reaction). SSO Advanced or iTaq Universal SYBR Green supermixes (Biorad) were used for qRT-PCR reactions in 96 well plates (25 ng cDNA per well), and Biorad CFX96 touch RT-PCR detection system was used. The ΔΔCT method was used for quantification, and the geometric mean of the CT values of three housekeeping genes (RPL8, RPL38, PSMA1) was used for normalization. The following primers were used:

*KLK3:* For (5′-CTTACCACCTGCACCCGGAG-3′) Rev (5′-TGCAGCACCAATCCACGTCA-3′)
*KLK2:* For (5′-AGAGGAGTTCTTGCGCCCC-3′) Rev (5′-CCCAGCACACAACATGAACTCT-3′)
*TMPRSS2:* For (5′-CCTCTGACTTTCAACGACCTAGTG-3′) Rev (5′-TCTTCCCTTTCTCCTCGGTGG-3′)
*NKX3-1:* For (5′-CAGAGACCGAGCCAGAAAGG-3′) Rev (5′-ACTCGATCACCTGAGTGTGGG-3′)
*AR:* For (5′-CCAGGGACCATGTTTTGCC-3′) Rev (5′-CGAAGACGACAAGATGGACAA-3′)
*AR Full:* For (5′-AGACAACCCAGAAGCTGACAGTG-3′) Rev (5′-GTGTAAGTTGCGGAAGCCAGG-3′)
*AR-V7:* For (5′-AATTGTCCATCTTGTCGTCTTCGG-3′)
  Rev (5′-GAGTCAGCAATCAAGAGAGTAGCC-3′)
*CDC25A:* For (5′-CATGAGAACTACAAACCTTGACAACC-3′)
  Rev (5′-CCCAGACATGCTCTTCCTCCTC-3′)
*EXO1:* For (5′-CTGCAGAGTTCAAATGCATCA-3′) Rev (5′-CGTAGCTTGGAGGTCTGGTC-3′)
*CENPN:* For (5′-TACACCGCTTCTGGGTCAGG-3′) Rev (5′-CTGTAGAGGTGTCGTAGAGTTGTGAG-3′)
*CDC45:* For (5′-CATGACAGCCTGTGCAACAC-3′) Rev (5′-GGGAAGACCCATGTCTGCAA-3′)
*SENP1:* For (5′-TTAGTACAGCAGAAGAGACAGTTCAAG-3′)
  Rev (5′-ACTGGAACTAAGACATCGAGACAGG-3′)
*CDK6:* For (5′-GATGGCTCTAACCTCAGTGGTCG-3′)
  Rev (5′-AGTTGATCAACATCTGAACTTCCACG-3′)
*CCND1:* For (5′-ATGCCAACCTCCTCAACGACC-3′) Rev (5′-CTGTTCCTCGCAGACCTCCAG-3′)
*RNF6:* For (5′-GAGAGATGGAACGAATTACAGAGACTC-3′)
  Rev (5′-CCAAACTAAACCGAAACTCTCCATTG-3′)
*PPP1CC:* For (5′-GGAGACGATCTGCCTCTTACTGG-3′)
  Rev (5′-TGAAGATCTGGTGATAAACCTCCATG-3′)
*EP300:* For (5′-CCAGATGGGAGGACAAACAGGA-3′) Rev (5′-CTGGCTGTTGACCCATGTTGG-3′)
*HOXB13:* For (5′-GGAAGGCAGCATTTGCAGACTC-3′) Rev (5′-CGCCTCTTGTCCTTGGTGATG-3′) *KMT2A:* For (5′-ATGGTGATGACAGTGCTAATGATGC-3′)
  Rev (5′-GTTGCTGGTGCAGGATGTGAGAC-3′)
*RPL8:* For (5′-CACCGTTATCTCCCACAACCCT-3′) Rev (5′-AGCCACCACACCAACCACAG-3′)
*RPL38:* For (5′-ACTTCCTGCTCACAGCCCGA-3′) Rev (5′-TCAGTTCCTTCACTGCCAAACCG-3′)
*PSMA1:* For (5′-CTGCCTGTGTCTCGTCTTGTATC-3′) Rev (5′-GGCCCATATCATCATAACCAGCA-3′)

Statistics: two-tailed t tests were used to calculate significant differential gene expression. n= 3. p <0.05 was considered to be statistically significant (FDR (q) was calculated when indicated to correct for multiple comparisons, q < 0.05 was considered significant).

### RNA-seq

LNCaP, LNCaP abl and 22Rv1 -/+ DHT cells were transfected with siGUGUGUA, siUGUUUGC, siNT-1, or siAR pool. Three repeats were conducted per siRNA in each cell line. RNA was extracted as for qRT-PCR at 40 hours, 48 hours and 64 hours after siRNA transfections in LNCaP, LNCaP abl and 22Rv1 -/+DHT cells, respectively. RNA integrity was ensured using Agilent 2100, and mRNAs were isolated using polyT coated beads and subsequently fragmented. Deoxythymidine triphosphate (dTTP) to deoxyuridine triphosphate (dUTP) containing cDNA libraries were generated, ligated to NEBNext Adaptor, PCR amplified and purified using AMPure XP beads. Sequencing was carried out with the Illumina Next-Generation sequencer at a depth of 20 million reads per sample. FastQC (Novogene) was used for quality control and the elimination of low-quality reads. STAR was used to map reads to the human genome/transcriptome and the DESeq2 R package was used to calculate differential gene expression. The Benjamini and Hochberg method was used for p-value adjustment, and average absolute log2 fold change > 0.5 relative to siNT transfected cells and adjusted p < 0.05 were used as cut offs for differential gene expression.

### Enrichment analyses

GSEA software version 4.2.3 was used for GSEAs^56,57^ of Hallmark gene sets (50 gene sets) from the Molecular Signature Database (MSigDB) (http://www.gsea-msigdb.org/gsea/index.jsp), as well as, AR regulated and predicted target gene sets. AR regulated gene sets were as follows: siAR up and down gene sets were generated from the top 150 significantly upregulated and downregulated genes, respectively, for each cell line; siAR-V7 up and down gene sets were obtained from Chen et al^32^ (GSE99378) and were selected by absolute log2 fold change > 1 and p-adjust < 0.05; LNCaP DHT up and down gene sets were obtained from Corbin et al^17^ (GSE165249) and were selected by the top 150 significantly upregulated and downregulated genes; AR Score was also utilized^30,31^. For the analyses, genes were rank ordered from upregulated to downregulated for each comparison relative to siNT transfected cells using the signal-to-noise ratio. 1,000 gene set permutations were used for each analysis and FDR < 0.1 was considered significant enrichment.

Essential gene lists were obtained from Fei et al (999 genes;^26^) and Blomen et al and Wang et al (1210 genes;^18,27,28^), and the AR coregulatory gene list was obtained from DePriest et al (274 genes;^25^). P-values for enrichment among downregulated genes were calculated by hypergeometric distribution and q-values were calculated to correct for multiple comparisons:

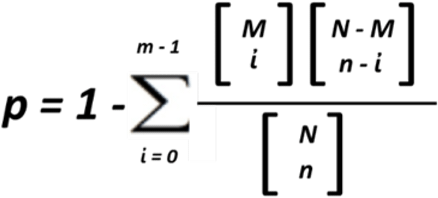

p = p value; N = number of total genes; M = number of genes in gene set; n = number of differentially expressed genes; i = number of overlapped genes of M and n.

### cWords and 3’UTR sequence analyses

Genes were rank ordered from downregulated to upregulated by signal-to-noise metric for each comparison based on the RNA seq data, and cWords^29^ webserver (http://servers.binf.ku.dk/cwords/) was used to determine 6 or 7 nucleotide 3’UTR sequences most associated with downregulation. Ensembl 103 MANE select transcript 3’UTR sequences were used for analysis. The downregulation of genes containing 3’UTR sequences complementary to the siRNA seeds was confirmed by Kolmogorov–Smirnov (KS) test, p < 0.05 was considered significant.

AR coregulatory and LNCaP essential gene sets were stratified by significant downregulation compared to siNT transfected cells (yes or no) and by the presence of 7mer and/or multiple 6mer seed match sequences in the 3’UTR. Statistics: Significant associations between downregulation and 3’UTR seed match sequences were determined by fisher’s exact test, p < 0.05 was considered significant.

## ACKNOWLEDGEMENTS

We are grateful to Dr. Zoran Culig (University of Innsbruck), Dr. Jun Luo (Johns Hopkins University) and Dr. S. Hayward (NorthShore Research Institute) for sharing cell lines. We appreciate Cody B. Bullock’s help with testing growth and viability of benign prostate cell lines. This work was supported by the Oklahoma IDeA Network of Biomedical Research Excellence (OK-INBRE; P20 GM103447; M.J.R.E.); and the National Institutes of Health (NIH 5U54GM104938; J.D.W.). Research reported in this publication was supported in part by the National Cancer Institute Cancer Center Support Grant (P30CA225520, COBRE P20GM103639), and the Oklahoma Tobacco Settlement Endowment Trust contract awarded to the University of Oklahoma Stephenson Cancer Center, and used the Tissue Pathology and the Molecular Biology and Cytometry Research Shared Resources. The content is solely the responsibility of the authors and does not necessarily represent the official views of the National Institutes of Health or the Oklahoma Tobacco Settlement Endowment Trust.

## AUTHOR CONTRIBUTIONS

Conceptualization, J.M.C. and M.J.R.E.; methodology, J.M.C. and M.J.R.E.; validation, J.M.C. and M.J.R.E.; formal analysis, J.M.C., L.W., C.G., J.D.W., C.X. and M.J.R.E.; investigation, J.M.C. and M.J.R.E.; resources, C.G., J.D.W., C.X. and M.B.; writing – original draft, J.M.C. and M.J.R.E.; writing – review & editing, J.M.C., C.G., J.D.W., C.X., M.B., A.S.A., and M.J.R.E; supervision, M.B., J.D.W., M.J.R.E.; funding acquisition, J.D.W. and M.J.R.E.

## REFERENCES

1. Siegel, R.L., Miller, K.D., Fuchs, H.E. and Jemal, A., Cancer statistics, 2022. CA Cancer J Clin, 2022. 72(1): p. 7–33.

2. Davey, R.A. and Grossmann, M., Androgen Receptor Structure, Function and Biology: From Bench to Bedside. Clin Biochem Rev, 2016. 37(1): p. 3–15.

3. Yuan, X., Cai, C., Chen, S., Chen, S., Yu, Z. and Balk, S.P., Androgen receptor functions in castration-resistant prostate cancer and mechanisms of resistance to new agents targeting the androgen axis. Oncogene, 2014. 33(22): p. 2815–25.

4. Huang, Y., Jiang, X., Liang, X. and Jiang, G., Molecular and cellular mechanisms of castration resistant prostate cancer. Oncol Lett, 2018. 15(5): p. 6063–6076.

5. Sharp, A., Welti, J., Blagg, J. and de Bono, J.S., Targeting Androgen Receptor Aberrations in Castration-Resistant Prostate Cancer. Clin Cancer Res, 2016. 22(17): p. 4280–2.

6. Bluemn, E.G., Coleman, I.M., Lucas, J.M., Coleman, R.T., Hernandez-Lopez, S., Tharakan, R., Bianchi-Frias, D., Dumpit, R.F., Kaipainen, A., Corella, A.N., et al., Androgen Receptor Pathway-Independent Prostate Cancer Is Sustained through FGF-Signaling. Cancer Cell, 2017. 32(4): p. 474–489 e6.

7. Coutinho, I., Day, T.K., Tilley, W.D. and Selth, L.A., Androgen receptor signaling in castration-resistant prostate cancer: a lesson in persistence. Endocr Relat Cancer, 2016. 23(12): p. T179–T197.

8. Jackson, A.L., Burchard, J., Schelter, J., Chau, B.N., Cleary, M., Lim, L. and Linsley, P.S., Widespread siRNA “off-target” transcript silencing mediated by seed region sequence complementarity. RNA, 2006. 12(7): p. 1179–87.

9. Zamore, P.D., Tuschl, T., Sharp, P.A. and Bartel, D.P., RNAi: double-stranded RNA directs the ATP-dependent cleavage of mRNA at 21 to 23 nucleotide intervals. Cell, 2000. 101(1): p. 25–33.

10. Peters, L. and Meister, G., Argonaute proteins: mediators of RNA silencing. Mol Cell, 2007. 26(5): p. 611–23.

11. Kehl, T., Backes, C., Kern, F., Fehlmann, T., Ludwig, N., Meese, E., Lenhof, H.P. and Keller, A., About miRNAs, miRNA seeds, target genes and target pathways. Oncotarget, 2017. 8(63): p. 107167–107175.

12. Bartel, D.P., MicroRNAs: target recognition and regulatory functions. Cell, 2009. 136(2): p. 215–33.

13. Lewis, B.P., Shih, I.H., Jones-Rhoades, M.W., Bartel, D.P. and Burge, C.B., Prediction of mammalian microRNA targets. Cell, 2003. 115(7): p. 787–98.

14. Fang, Z. and Rajewsky, N., The impact of miRNA target sites in coding sequences and in 3’UTRs. PLoS One, 2011. 6(3): p. e18067.

15. Satoh, J. and Tabunoki, H., Comprehensive analysis of human microRNA target networks. BioData Min, 2011. 4: p. 17.

16. Ebert, M.S. and Sharp, P.A., Roles for microRNAs in conferring robustness to biological processes. Cell, 2012. 149(3): p. 515–24.

17. Corbin, J.M., Georgescu, C., Wren, J.D., Xu, C., Asch, A.S. and Ruiz-Echevarria, M.J., Seed-mediated RNA interference of androgen signaling and survival networks induces cell death in prostate cancer cells. Mol Ther Nucleic Acids, 2021. 24: p. 337–351.

18. Putzbach, W., Gao, Q.Q., Patel, M., van Dongen, S., Haluck-Kangas, A., Sarshad, A.A., Bartom, E.T., Kim, K.A., Scholtens, D.M., Hafner, M., et al., Many si/shRNAs can kill cancer cells by targeting multiple survival genes through an off-target mechanism. Elife, 2017. **6**.

19. Gao, Q.Q., Putzbach, W.E., Murmann, A.E., Chen, S., Sarshad, A.A., Peter, J.M., Bartom, E.T., Hafner, M. and Peter, M.E., 6mer seed toxicity in tumor suppressive microRNAs. Nat Commun, 2018. 9(1): p. 4504.

20. Murmann, A.E., McMahon, K.M., Haluck-Kangas, A., Ravindran, N., Patel, M., Law, C.Y., Brockway, S., Wei, J.J., Thaxton, C.S. and Peter, M.E., Induction of DISE in ovarian cancer cells in vivo. Oncotarget, 2017. 8(49): p. 84643–84658.

21. Li, Q.V., Dixon, G., Verma, N., Rosen, B.P., Gordillo, M., Luo, R., Xu, C., Wang, Q., Soh, C.L., Yang, D., et al., Genome-scale screens identify JNK-JUN signaling as a barrier for pluripotency exit and endoderm differentiation. Nat Genet, 2019. 51(6): p. 999–1010.

22. Zhang, L., Liao, Y. and Tang, L., MicroRNA-34 family: a potential tumor suppressor and therapeutic candidate in cancer. J Exp Clin Cancer Res, 2019. 38(1): p. 53.

23. Hagman, Z., Larne, O., Edsjo, A., Bjartell, A., Ehrnstrom, R.A., Ulmert, D., Lilja, H. and Ceder, Y., miR-34c is downregulated in prostate cancer and exerts tumor suppressive functions. Int J Cancer, 2010. 127(12): p. 2768–76.

24. Ostling, P., Leivonen, S.K., Aakula, A., Kohonen, P., Makela, R., Hagman, Z., Edsjo, A., Kangaspeska, S., Edgren, H., Nicorici, D., et al., Systematic analysis of microRNAs targeting the androgen receptor in prostate cancer cells. Cancer Res, 2011. 71(5): p. 1956–67.

25. DePriest, A.D., Fiandalo, M.V., Schlanger, S., Heemers, F., Mohler, J.L., Liu, S. and Heemers, H.V., Regulators of Androgen Action Resource: a one-stop shop for the comprehensive study of androgen receptor action. Database (Oxford), 2016. **2016**.

26. Fei, T., Chen, Y., Xiao, T., Li, W., Cato, L., Zhang, P., Cotter, M.B., Bowden, M., Lis, R.T., Zhao, S.G., et al., Genome-wide CRISPR screen identifies HNRNPL as a prostate cancer dependency regulating RNA splicing. Proc Natl Acad Sci U S A, 2017. 114(26): p. E5207–E5215.

27. Blomen, V.A., Majek, P., Jae, L.T., Bigenzahn, J.W., Nieuwenhuis, J., Staring, J., Sacco, R., van Diemen, F.R., Olk, N., Stukalov, A., et al., Gene essentiality and synthetic lethality in haploid human cells. Science, 2015. 350(6264): p. 1092–6.

28. Wang, T., Birsoy, K., Hughes, N.W., Krupczak, K.M., Post, Y., Wei, J.J., Lander, E.S. and Sabatini, D.M., Identification and characterization of essential genes in the human genome. Science, 2015. 350(6264): p. 1096–101.

29. Rasmussen, S.H., Jacobsen, A. and Krogh, A., cWords - systematic microRNA regulatory motif discovery from mRNA expression data. Silence, 2013. 4(1): p. 2.

30. Hieronymus, H., Lamb, J., Ross, K.N., Peng, X.P., Clement, C., Rodina, A., Nieto, M., Du, J., Stegmaier, K., Raj, S.M., et al., Gene expression signature-based chemical genomic prediction identifies a novel class of HSP90 pathway modulators. Cancer Cell, 2006. 10(4): p. 321–30.

31. Hu, C., Fang, D., Xu, H., Wang, Q. and Xia, H., The androgen receptor expression and association with patient’s survival in different cancers. Genomics, 2020. 112(2): p. 1926–1940.

32. Chen, Z., Wu, D., Thomas-Ahner, J.M., Lu, C., Zhao, P., Zhang, Q., Geraghty, C., Yan, P.S., Hankey, W., Sunkel, B., et al., Diverse AR-V7 cistromes in castration-resistant prostate cancer are governed by HoxB13. Proc Natl Acad Sci U S A, 2018. 115(26): p. 6810–6815.

33. Hadji, A., Ceppi, P., Murmann, A.E., Brockway, S., Pattanayak, A., Bhinder, B., Hau, A., De Chant, S., Parimi, V., Kolesza, P., et al., Death induced by CD95 or CD95 ligand elimination. Cell Rep, 2014. 7(1): p. 208–22.

34. Powers, E., Karachaliou, G.S., Kao, C., Harrison, M.R., Hoimes, C.J., George, D.J., Armstrong, A.J. and Zhang, T., Novel therapies are changing treatment paradigms in metastatic prostate cancer. J Hematol Oncol, 2020. 13(1): p. 144.

35. Saad, F., Bogemann, M., Suzuki, K. and Shore, N., Treatment of nonmetastatic castration-resistant prostate cancer: focus on second-generation androgen receptor inhibitors. Prostate Cancer Prostatic Dis, 2021. 24(2): p. 323–334.

36. Clarke, N., Wiechno, P., Alekseev, B., Sala, N., Jones, R., Kocak, I., Chiuri, V.E., Jassem, J., Flechon, A., Redfern, C., et al., Olaparib combined with abiraterone in patients with metastatic castration-resistant prostate cancer: a randomised, double-blind, placebo-controlled, phase 2 trial. Lancet Oncol, 2018. 19(7): p. 975–986.

37. Attard, G., Murphy, L., Clarke, N.W., Cross, W., Jones, R.J., Parker, C.C., Gillessen, S., Cook, A., Brawley, C., Amos, C.L., et al., Abiraterone acetate and prednisolone with or without enzalutamide for high-risk non-metastatic prostate cancer: a meta-analysis of primary results from two randomised controlled phase 3 trials of the STAMPEDE platform protocol. Lancet, 2022. 399(10323): p. 447–460.

38. Sharma, P., Pachynski, R.K., Narayan, V., Flechon, A., Gravis, G., Galsky, M.D., Mahammedi, H., Patnaik, A., Subudhi, S.K., Ciprotti, M., et al., Nivolumab Plus Ipilimumab for Metastatic Castration-Resistant Prostate Cancer: Preliminary Analysis of Patients in the CheckMate 650 Trial. Cancer Cell, 2020. 38(4): p. 489–499 e3.

39. Shenderov, E., Boudadi, K., Fu, W., Wang, H., Sullivan, R., Jordan, A., Dowling, D., Harb, R., Schonhoft, J., Jendrisak, A., et al., Nivolumab plus ipilimumab, with or without enzalutamide, in AR-V7-expressing metastatic castration-resistant prostate cancer: A phase-2 nonrandomized clinical trial. Prostate, 2021. 81(6): p. 326–338.

40. Jackson, A.L., Burchard, J., Leake, D., Reynolds, A., Schelter, J., Guo, J., Johnson, J.M., Lim, L., Karpilow, J., Nichols, K., et al., Position-specific chemical modification of siRNAs reduces “off-target” transcript silencing. RNA, 2006. 12(7): p. 1197–205.

41. Jackson, A.L. and Linsley, P.S., Recognizing and avoiding siRNA off-target effects for target identification and therapeutic application. Nat Rev Drug Discov, 2010. 9(1): p. 57–67.

42. Kamola, P.J., Nakano, Y., Takahashi, T., Wilson, P.A. and Ui-Tei, K., The siRNA Non-seed Region and Its Target Sequences Are Auxiliary Determinants of Off-Target Effects. PLoS Comput Biol, 2015. 11(12): p. e1004656.

43. Fletcher, C.E., Sulpice, E., Combe, S., Shibakawa, A., Leach, D.A., Hamilton, M.P., Chrysostomou, S.L., Sharp, A., Welti, J., Yuan, W., et al., Androgen receptor-modulatory microRNAs provide insight into therapy resistance and therapeutic targets in advanced prostate cancer. Oncogene, 2019. 38(28): p. 5700–5724.

44. Bielska, A., Skwarska, A., Kretowski, A. and Niemira, M., The Role of Androgen Receptor and microRNA Interactions in Androgen-Dependent Diseases. Int J Mol Sci, 2022. 23(3).

45. Coarfa, C., Fiskus, W., Eedunuri, V.K., Rajapakshe, K., Foley, C., Chew, S.A., Shah, S.S., Geng, C., Shou, J., Mohamed, J.S., et al., Comprehensive proteomic profiling identifies the androgen receptor axis and other signaling pathways as targets of microRNAs suppressed in metastatic prostate cancer. Oncogene, 2016. 35(18): p. 2345–56.

46. Meng, Z. and Lu, M., RNA Interference-Induced Innate Immunity, Off-Target Effect, or Immune Adjuvant? Front Immunol, 2017. 8: p. 331.

47. Hornung, V., Guenthner-Biller, M., Bourquin, C., Ablasser, A., Schlee, M., Uematsu, S., Noronha, A., Manoharan, M., Akira, S., de Fougerolles, A., et al., Sequence-specific potent induction of IFN-alpha by short interfering RNA in plasmacytoid dendritic cells through TLR7. Nat Med, 2005. 11(3): p. 263–70.

48. Forsbach, A., Nemorin, J.G., Montino, C., Muller, C., Samulowitz, U., Vicari, A.P., Jurk, M., Mutwiri, G.K., Krieg, A.M., Lipford, G.B., et al., Identification of RNA sequence motifs stimulating sequence-specific TLR8-dependent immune responses. J Immunol, 2008. 180(6): p. 3729–38.

49. Bartoszewski, R. and Sikorski, A.F., Editorial focus: understanding off-target effects as the key to successful RNAi therapy. Cell Mol Biol Lett, 2019. 24: p. 69.

50. Sioud, M., Single-stranded small interfering RNA are more immunostimulatory than their double-stranded counterparts: a central role for 2′-hydroxyl uridines in immune responses. Eur J Immunol, 2006. 36(5): p. 1222–30.

51. Sajid, M.I., Moazzam, M., Kato, S., Yeseom Cho, K. and Tiwari, R.K., Overcoming Barriers for siRNA Therapeutics: From Bench to Bedside. Pharmaceuticals (Basel), 2020. 13(10).

52. Lee, Y., Urban, J.H., Xu, L., Sullenger, B.A. and Lee, J., 2′Fluoro Modification Differentially Modulates the Ability of RNAs to Activate Pattern Recognition Receptors. Nucleic Acid Ther, 2016. 26(3): p. 173–82.

53. Deleavey, G.F., Watts, J.K. and Damha, M.J., Chemical modification of siRNA. Curr Protoc Nucleic Acid Chem, 2009. **Chapter 16**: p. Unit 16 3.

54. Jiang, M., Strand, D.W., Fernandez, S., He, Y., Yi, Y., Birbach, A., Qiu, Q., Schmid, J., Tang, D.G. and Hayward, S.W., Functional remodeling of benign human prostatic tissues in vivo by spontaneously immortalized progenitor and intermediate cells. Stem Cells, 2010. 28(2): p. 344–56.

55. Gu, S., Jin, L., Zhang, Y., Huang, Y., Zhang, F., Valdmanis, P.N. and Kay, M.A., The loop position of shRNAs and pre-miRNAs is critical for the accuracy of dicer processing in vivo. Cell, 2012. 151(4): p. 900–911.

56. Mootha, V.K., Lindgren, C.M., Eriksson, K.F., Subramanian, A., Sihag, S., Lehar, J., Puigserver, P., Carlsson, E., Ridderstrale, M., Laurila, E., et al., PGC-1alpha-responsive genes involved in oxidative phosphorylation are coordinately downregulated in human diabetes. Nat Genet, 2003. 34(3): p. 267–73.

57. Subramanian, A., Tamayo, P., Mootha, V.K., Mukherjee, S., Ebert, B.L., Gillette, M.A., Paulovich, A., Pomeroy, S.L., Golub, T.R., Lander, E.S., et al., Gene set enrichment analysis: a knowledge-based approach for interpreting genome-wide expression profiles. Proc Natl Acad Sci U S A, 2005. 102(43): p. 15545–50.

