## Supplementary material for "A NOVEL UNBIASED SEED-BASED RNAi SCREEN IDENTIFIES SMALL RNAs THAT INHIBIT ANDROGEN SIGNALING AND PROSTATE CANCER CELL GROWTH": Suppl. Figures and Table 1

**SUPPLEMENTAL FIGURES**

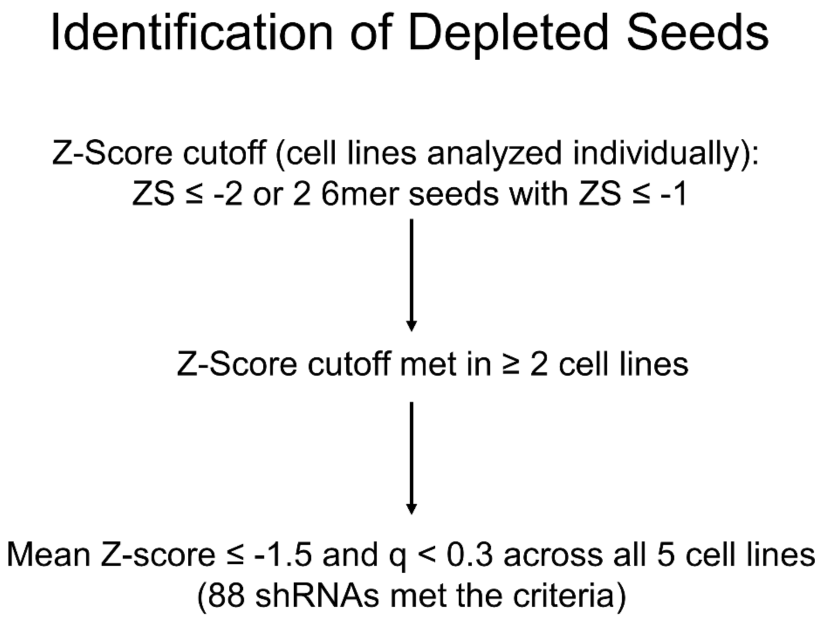


**Figure S1: Criteria for the identification of depleted shRNAs**

Z-scores were calculated for relative shRNA abundance comparing number of reads determined by NGS at ten doublings to number of reads at zero doublings, as previously described.[21] In individual cell lines, shRNAs with a z-score ≤ -2 or with two 6mer seeds within the 7mer seed with a z-score ≤ -1 met the initial cutoff. shRNAs that met the z-score cutoff in at least two cell lines and had a mean z-score across all cell lines ≤ -1.5 and $q < 0.3$ were considered to be depleted (FDR of all z-scores across the 5 cell lines).

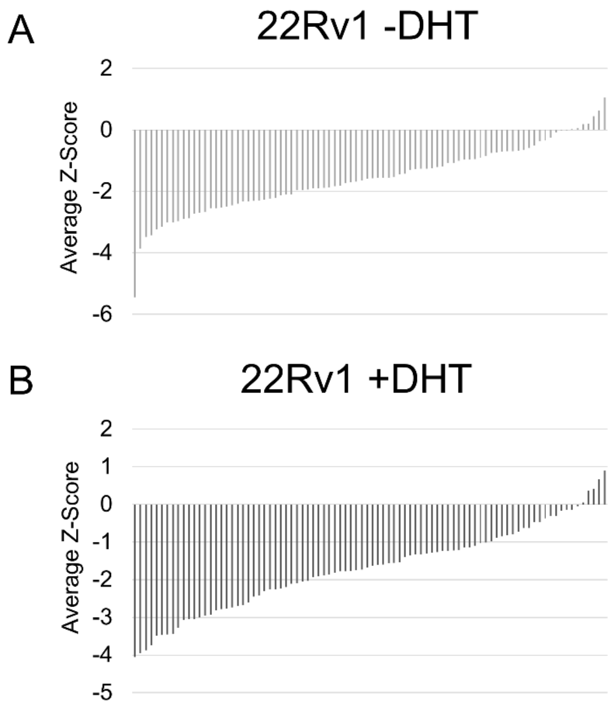


**Figure S2: The 88 negatively selected shRNAs were mostly depleted at the two doubling time point in 22Rv1 cells**

Bar graphs showing the average z-scores for the 88 shRNAs negatively selected across all cell lines at the two doubling time point in 22Rv1 cells grown in the absence (top) and presence (bottom) of DHT.

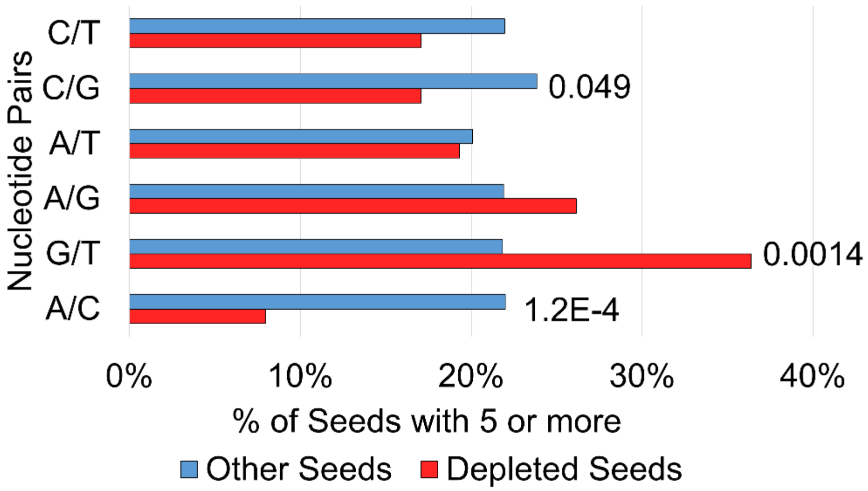


**Figure S3: Depleted shRNAs showed a tendency towards G/T rich seed sequences**

Bar graph showing the percentage of seeds with 5 or more of the indicated nucleotide pairs within the seed sequence for the group of depleted shRNAs (red) or the other shRNAs (blue). $P < 0.05$ was considered a significant association as calculated by hypergeometric distribution.

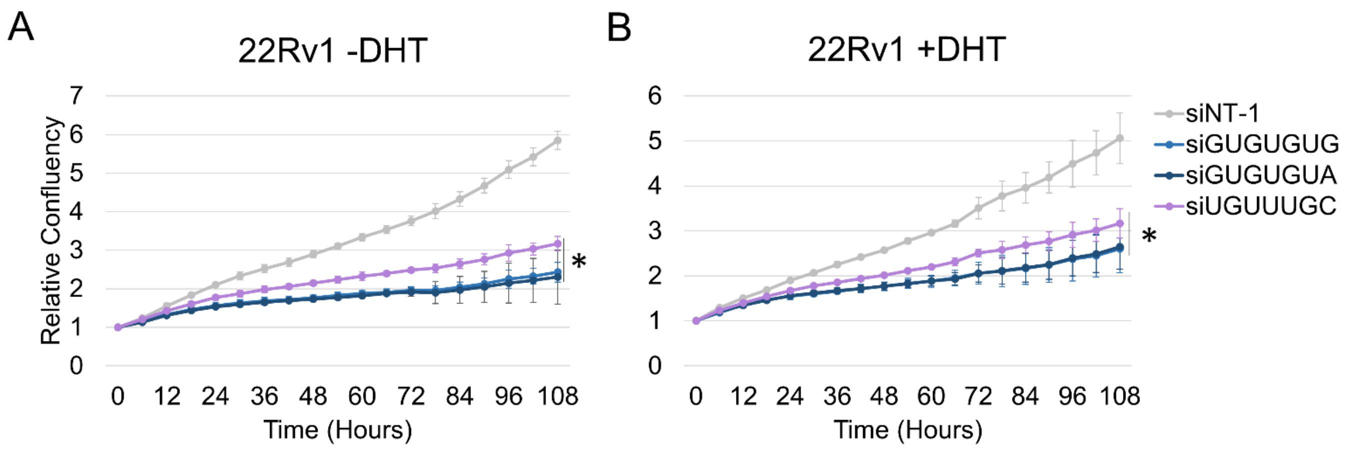


**Figure S4: siGUGUGUA and siUGUUUGC inhibit growth of 22Rv1 cells grown in the presence or absence of DHT**

Cell growth of 22Rv1 cells grown in the absence (A) or presence (B) of 1 nM DHT and transfected with the indicated siRNAs. Relative confluency, as measured by InCucyte, was used as a surrogate for cell growth. * $p < 0.05$ as determined by T-test, N= 3.

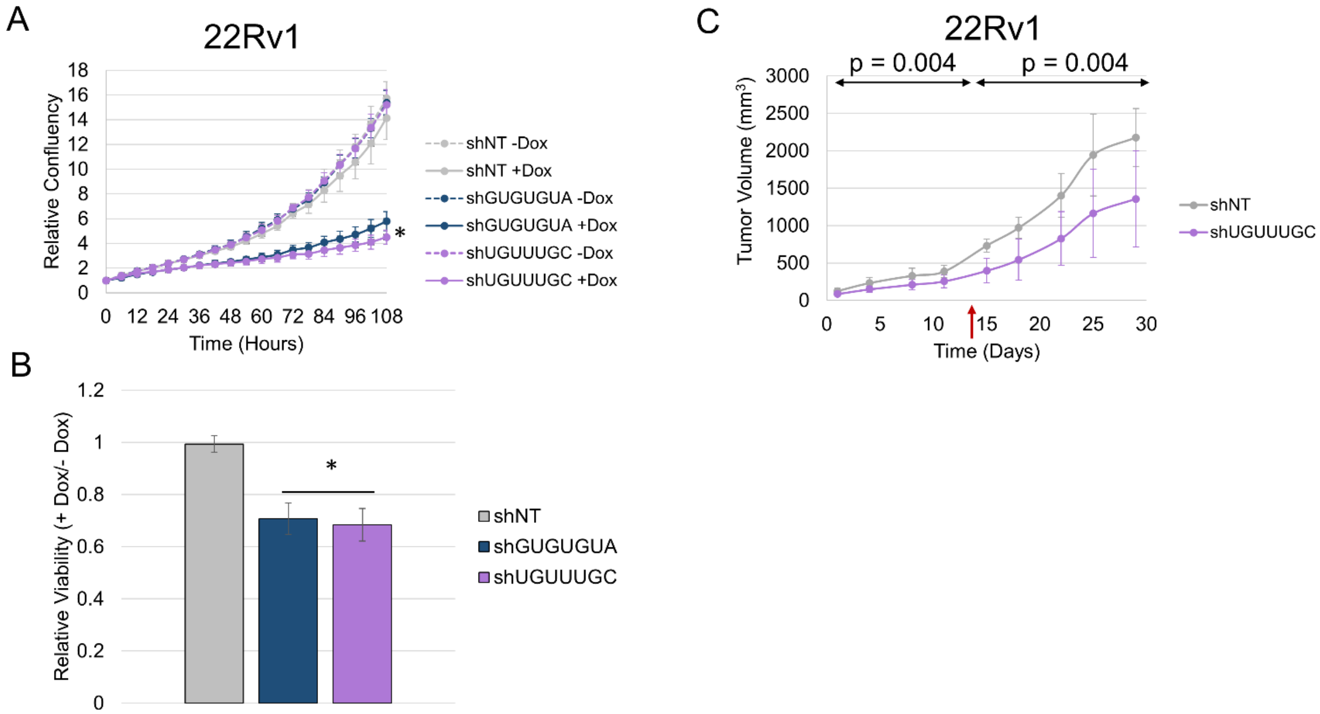


**Figure S5: shGUGUGUA and shUGUUUGC reduce 22Rv1 cell growth, viability and xenograft tumor growth**

22Rv1 shGUGUGUA, shUGUUUGC and shNT cell growth (A) and viability (B) in response to 500 ng/ml doxycycline. Relative confluency, as measured by InCucyte, was used as a surrogate for cell growth and relative viability was measured by trypan blue. * $p < 0.05$ as determined by T-test, N= 3. C) Plot showing 22Rv1 shUGUUUGC and shNT tumor growth in mouse xenograft models. NRG mice were pre-fed two days on doxycycline containing chow prior to subcutaneous injection of cells into the flanks. Tumor size was measured three times per week. Significance was determined using a segmented linear mixed model, and day 13 was identified as the transition day (red arrow). P values for before and after the transition day are labeled on the plot. n = 5 for shNT; n = 6 for shUGUUUGC.

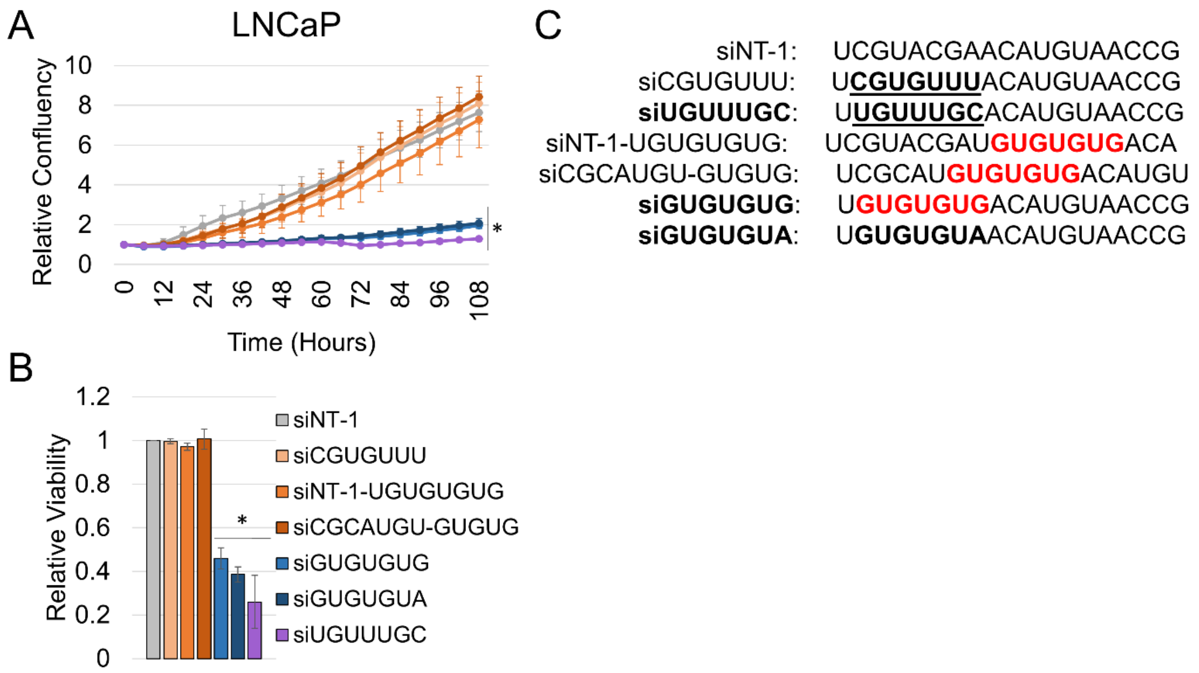


**Figure S6: Shifting or rearranging seed sequences within the siRNAs reverse the toxic effects**

LNCaP growth (A) and viability (B) in response to transfection with siRNAs (C) (Sequences correspond to the guide strand), including an siRNA with same nucleotide content as siUGUUUGC but rearranged order (underlined) and two siRNAs with the GUGUGUG sequence (red) shifted in the 3' direction. Relative confluency, as measured by InCucyte, was used as a surrogate for cell growth and relative viability was measured by trypan blue. * $q < 0.05$ compared to siNT-1 transfection was considered significant as determined by T-test, N= 3.

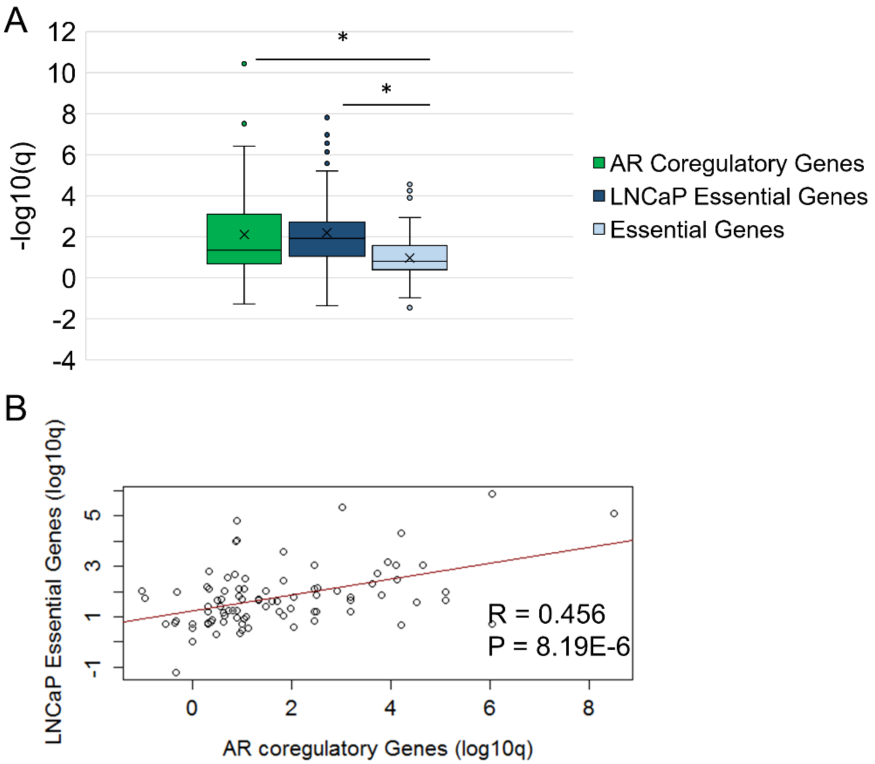


**Figure S7: AR coregulatory and LNCaP (PCa) essential genes are more significantly enriched among depleted shRNA predicted targets than non-PCa essential genes.**

A) Box and whisker plot comparing the enrichment (-log10(q)) of AR coregulatory[25], and LNCaP (PCa) essential[26] and non-PCa essential[18,27,28] gene sets among the predicted targets of the depleted shRNAs (n=88). *$p < 0.05$, as determined by t-test. B) Scatter plot showing the -log(q) of enrichment of AR coregulatory, LNCaP (PCa) essential gene sets among the predicted targets of the 88 depleted shRNAs. Each dot represents a single shRNA.

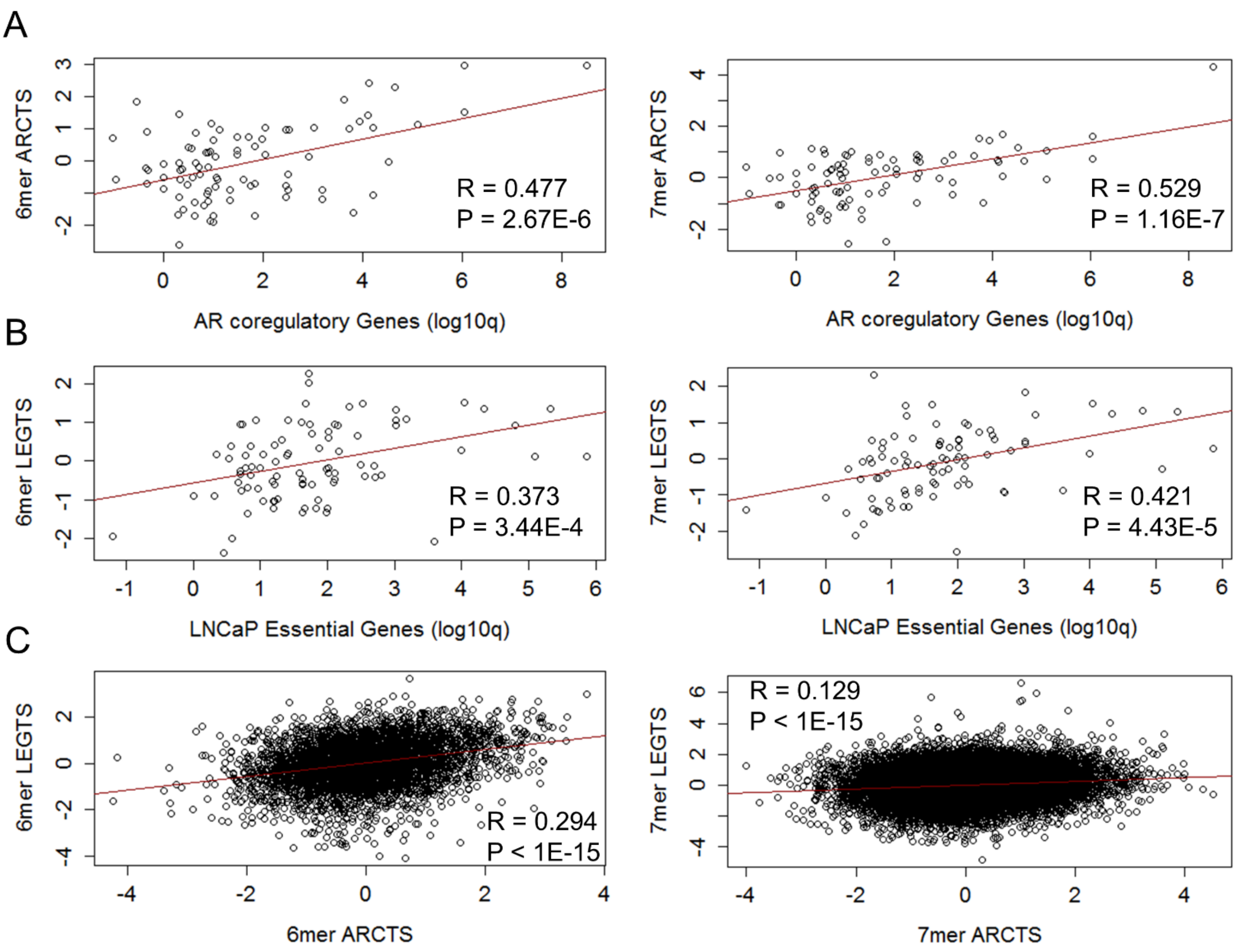


**Figure S8: Significant correlations between enrichment of gene sets from shRNA predicted targets and abundance of 6 and 7 nucleotide shRNA seed complementary sequences in the 3'UTR of AR coregulatory and essential genes**

(A-B) Scatter plots showing correlations between enrichment (-log10q) of AR coregulatory[25] (A) and LNCaP essential genes[26] (B) among the predicted targets of 88 depleted shRNAs and the relative abundance of 6 (Left) and 7 (Right) nucleotide 3'UTR sequences complementary to the seed sequences of the depleted shRNAs within AR Coregulatory (A, ARCTS) and LNCaP essential genes (B, LEGTS). Each dot represents a single shRNA. R and p values showing significant correlations are located on the plots. C) Scatter plots showing the correlation between the relative abundance of individual 6mer (Left) and 7mer (Right) seed matches from all screen shRNAs (n = 15,572) in the 3'UTR of AR coregulatory (ARCTS), LNCaP essential (LEGTS). Each dot represents a single shRNA. R and p values for significant correlations are labeled on the plots. ARCTS = AR Coregulatory Target Score, LEGTS = LNCaP Essential Gene Target Score. See materials and methods for a description of these scores.

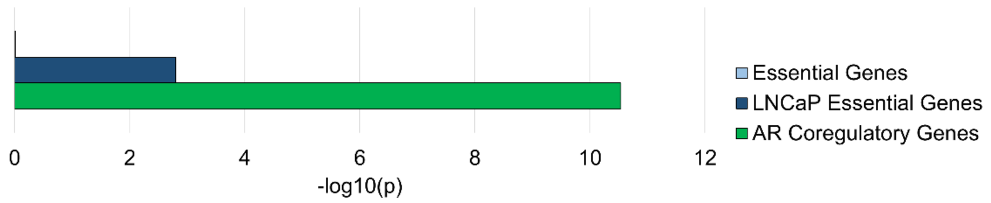


**Figure S9: AR coregulatory and LNCaP essential genes sets are enriched among genes that are common predicted targets of >9 depleted shRNAs**

Bar graph showing the significance (-log10p) of enrichment of AR coregulatory[25] and LNCaP essential[26] gene sets among genes predicted to be targeted by > 10% of depleted seeds. Significance was calculated by hypergeometric distribution.

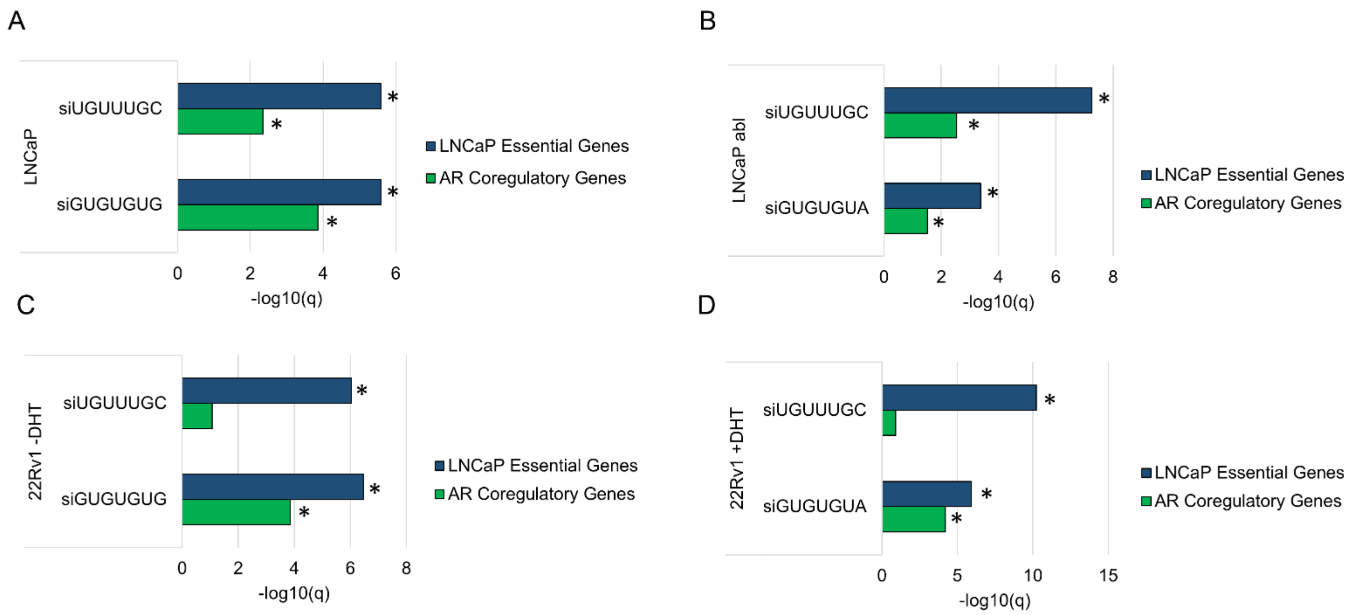


**Figure S10: AR coregulatory and LNCaP essential gene sets are enriched among the genes downregulated by siGUGUGUA and siUGUUUGC in PCa cell lines**

Bar graphs of –log10(q) of the enrichment of LNCaP essential[26] and AR coregulatory[25] gene sets among genes downregulated by siGUGUGUA and siUGUUUGC in LNCaP (A), LNCaP abl (B), 22Rv1 –DHT (C), and 22Rv1 +DHT (D) cells. q-values were calculated by hypergeometric distribution, * $q < 0.05$ or $–\log10(q) > 1.3$ was considered significant.

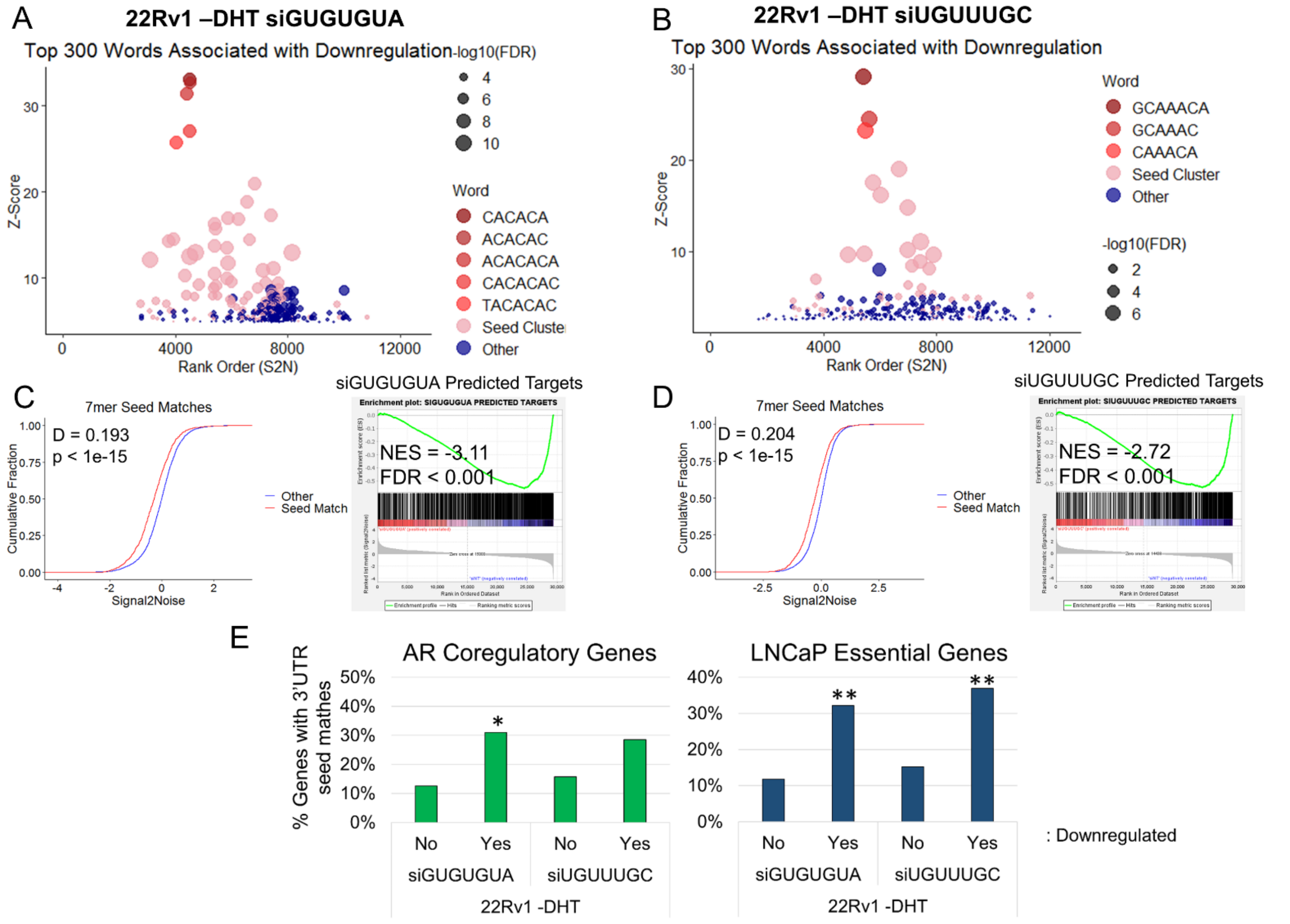


**Figure S11: AR coregulatory and prostate cancer essential gene downregulation in 22Rv1 –DHT cells is associated with sequences complementary to the GUGUGUA and UGUUUGC seeds in the 3'UTR**

(A-B) cWords plots displaying the top 300 words (6 and 7 nucleotide 3'UTR sequences) associated with gene downregulation in 22Rv1 –DHT cells transfected with siGUGUGUA (A) or siUGUUUGC (B). The top seed-related sequences are indicated; "seed cluster" includes other seed-related sequences. Genes were ranked ordered by signal to noise values from GSEA. (C-D) eCDF plots from 22Rv1 -DHT cells transfected with siGUGUGUA (C, left) or siUGUUUGC (D, left) showing the cumulative fraction of genes across the signal to noise rank ordered gene expression list, for genes with and without 7mer seed match sequences in their 3'UTR ; $p < 0.05$ as determined by KS test, D and p values are labeled on plots. GSEA enrichment plots for miRDB predicted target gene sets in 22Rv1 -DHT cells transfected with siGUGUGUA (C, right) or siUGUUUGC (D, right). E) Bar graphs showing the percentage of AR coregulatory[25] (Left) and LNCaP essential[26] (Right) genes with 7mer and/or multiple 6mer seed matches to siGUGUGUA or siUGUUUGC in the 3' UTR, stratified by whether the genes are downregulated (Yes) or not (No) by the corresponding siRNAs in 22Rv1 +DHT cells. *$p < 0.05$, **$p < 0.0001$, as determined by Fisher's exact test.

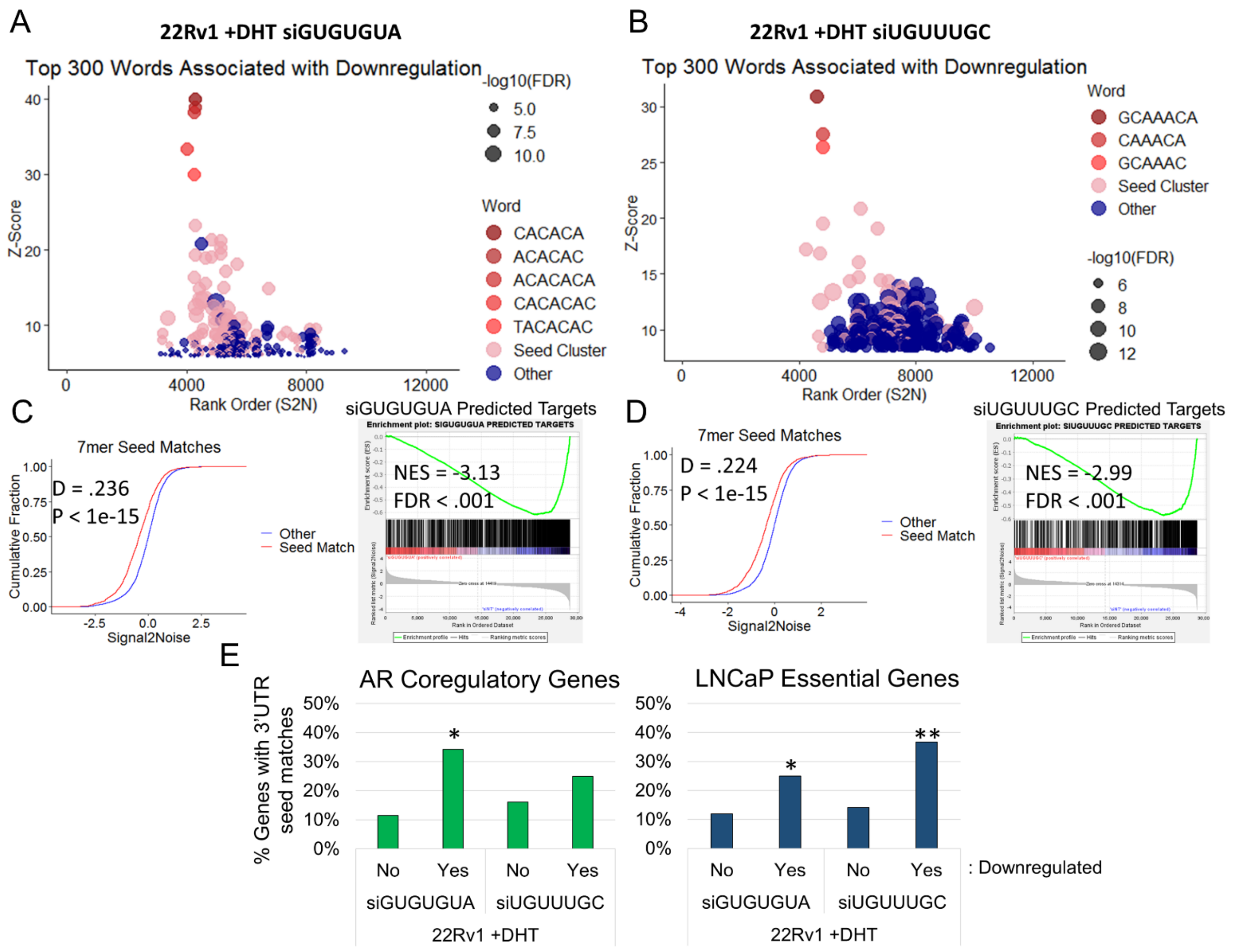


**Figure S12: Global, AR coregulatory and prostate cancer essential gene downregulation in 22Rv1 +DHT cells is associated with sequences complementary to the GUGUGUA and UGUUUGC seeds in the 3'UTR**

(A-B) cWords plots displaying the top 300 words (6 and 7 nucleotide 3'UTR sequences) associated with gene downregulation in 22Rv1 +DHT cells transfected with siGUGUGUA (A) or siUGUUUGC (B). The top seed-related sequences are indicated; "seed cluster" includes other seed-related sequences. Genes were ranked ordered by signal to noise values from GSEA. (C-D) eCDF plots from 22Rv1 +DHT cells transfected with siGUGUGUA (C, left) or siUGUUUGC (D, left) showing the cumulative fraction of genes across the signal to noise rank ordered gene expression list, for genes with and without 7mer seed match sequences in their 3'UTR ; $p < 0.05$ as determined by KS test, D and p values are labeled on plots. GSEA enrichment plots for miRDB predicted target gene sets in 22Rv1 +DHT cells transfected with siGUGUGUA (C, right) or siUGUUUGC (D, right). E) Bar graphs showing the percentage of AR coregulatory[25] (Left) and LNCaP essential[26] (Right) genes with 7mer and/or multiple 6mer seed matches to siGUGUGUA or siUGUUUGC in the 3' UTR, stratified by whether the genes are downregulated (Yes) or not (No) by the corresponding siRNAs in 22Rv1 +DHT cells. *$p < 0.05$, **$p < 0.0001$, as determined by Fisher's exact test.

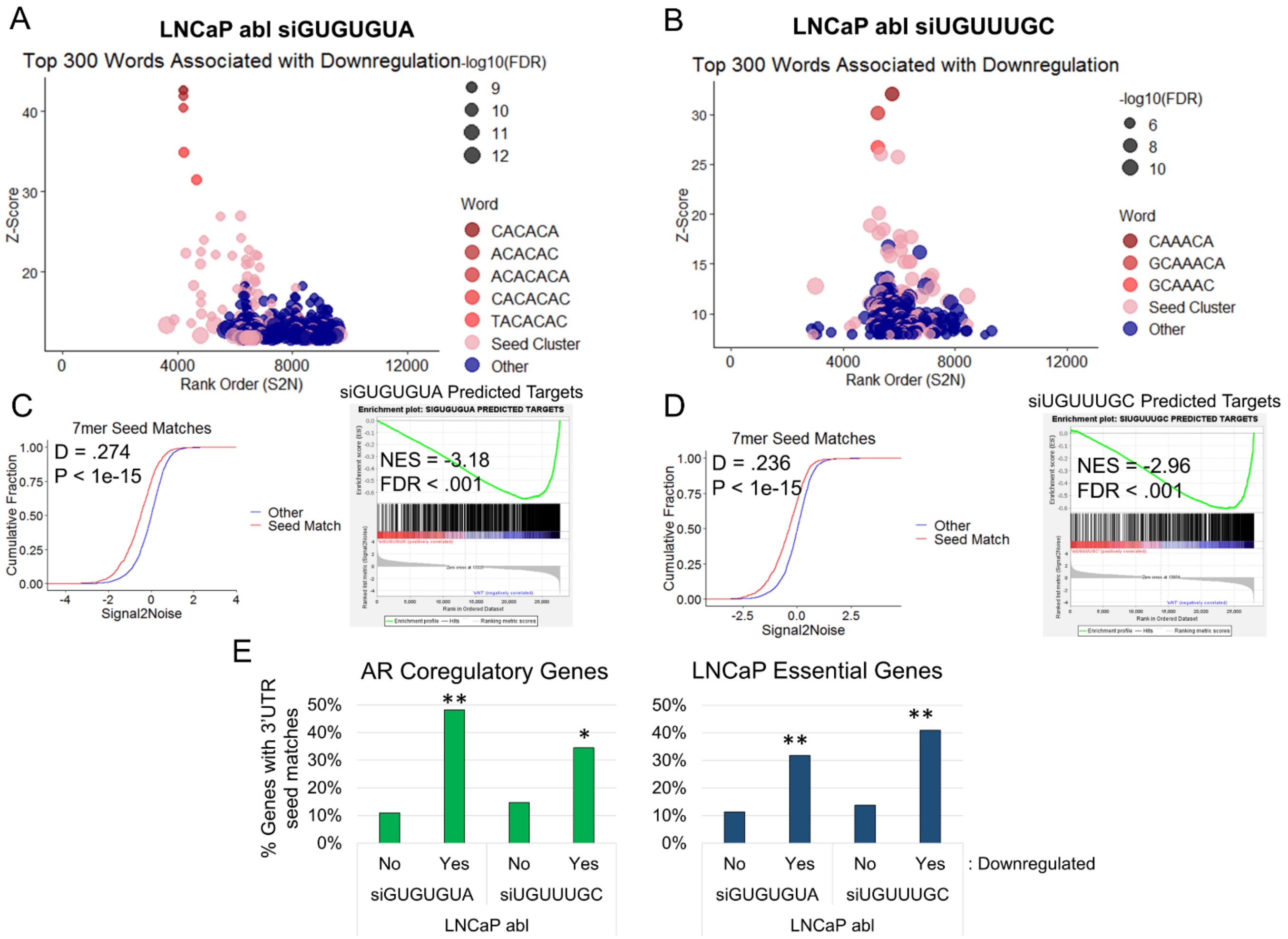


**Figure S13: Global, AR coregulatory and prostate cancer essential gene downregulation in LNCaP abl cells is associated with sequences complementary to the GUGUGUA and UGUUUGC seeds in the 3'UTR**

(A-B) cWords plots displaying the top 300 words (6 and 7 nucleotide 3'UTR sequences) associated with gene downregulation in LNCaP abl cells transfected with siGUGUGUA (A) or siUGUUUGC (B). The top seed-related sequences are indicated; "seed cluster" includes other seed-related sequences. Genes were ranked ordered by signal to noise values from GSEA. (C-D) eCDF plots from LNCaP abl cells transfected with siGUGUGUA (C, left) or siUGUUUGC (D, left) showing the cumulative fraction of genes across the signal to noise rank ordered gene expression list, for genes with and without 7mer seed match sequences in their 3'UTR ; $p < 0.05$ as determined by KS test, D and p values are labeled on plots. GSEA enrichment plots for miRDB predicted target gene sets in LNCaP abl cells transfected with siGUGUGUA (C, right) or siUGUUUGC (D, right). E) Bar graphs showing the percentage of AR coregulatory[25] (Left) and LNCaP[26] essential (Right) genes with 7mer and/or multiple 6mer seed matches to siGUGUGUA or siUGUUUGC in the 3' UTR, stratified by whether the genes are downregulated (Yes) or not (No) by the corresponding siRNAs in LNCaP abl cells. *$p < 0.05$, **$p < 0.0001$, as determined by Fisher's exact test.

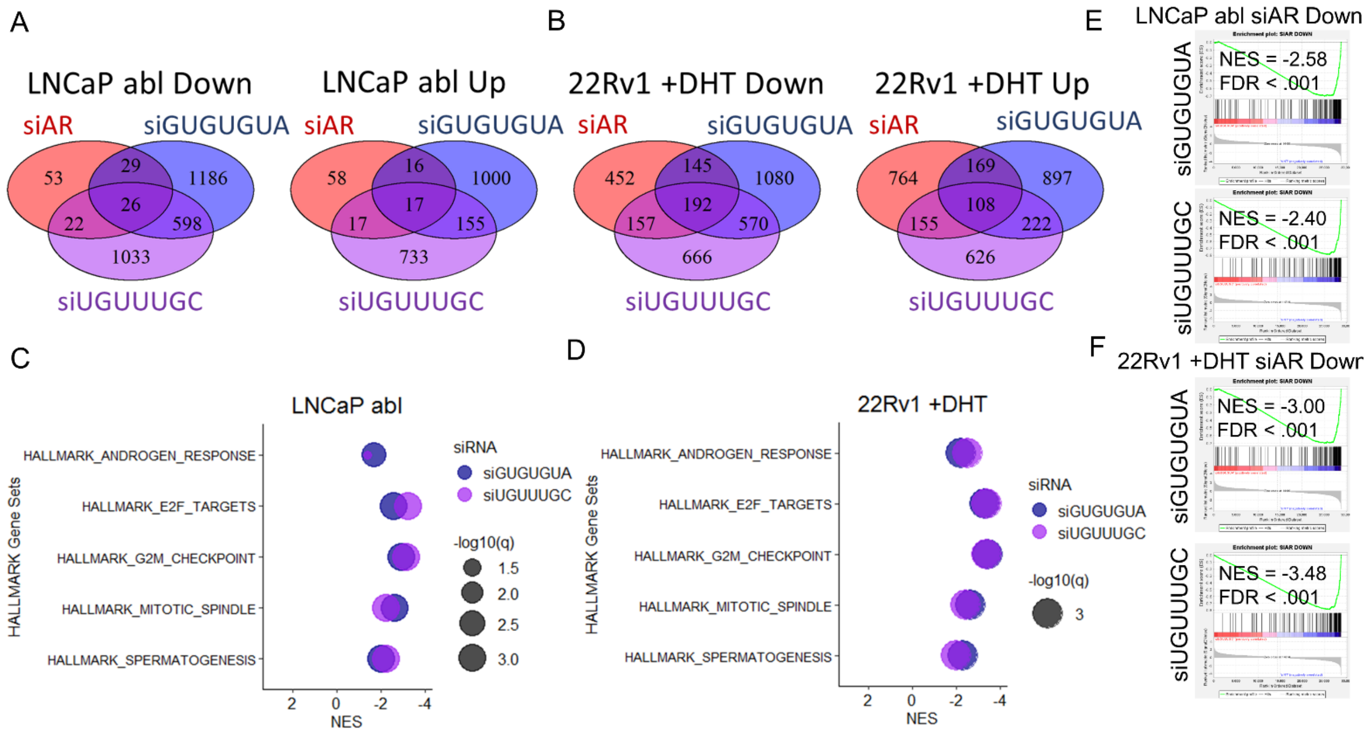


**Figure S14: GUGUGUA and UGUUUGC seed sequences reduce the expression of androgen responsive and cell cycle genes**

(A-B) Venn diagrams showing the overlap of significantly upregulated and downregulated genes as determined by RNA-seq ([log2(FC)] > 0.5, padj < 0.05) in LNCaP abl cells (A) and 22Rv1 cells grown in the presence of DHT (B) transfected with siGUGUGUA, siUGUUUGC or AR targeted siRNA (siAR), compared to cells transfected with siNT-1. (C-D) Plots showing NES and –log10(q) for significantly upregulated and downregulated Hallmark gene sets in response to transfection with siGUGUGUA and siUGUUUGC in LNCaP abl (C) and 22Rv1 +DHT (D) cells. (E-F) GSEA enrichment plots showing the enrichment of the “siAR down” gene set in LNCaP abl cells (E) and 22Rv1 +DHT cells (F) transfected with siGUGUGUA and siUGUUUGC. Positive and negative NES indicate upregulated and downregulated gene sets respectively. FDR < 0.1 was considered significant.

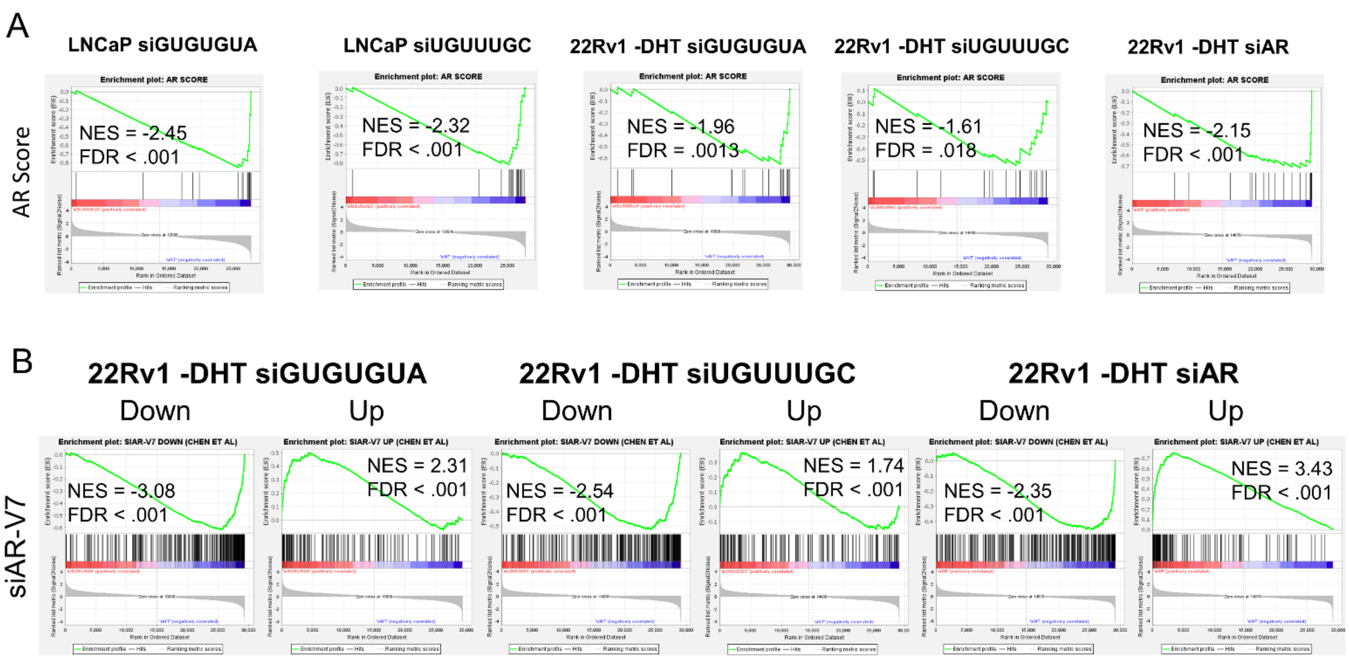


**Figure S15: AR regulated gene sets are downregulated by siGUGUGUA and siUGUUUGC in LNCaP and 22Rv1-DHT cells**

GSEA enrichment plots of "AR score"[30,31] (A) and "siAR-V7 up" and "siAR-V7 down"[32] (B) gene sets in LNCaP and 22Rv1 -DHT cells transfected with siGUGUGUA, siUGUUUGC or siAR.

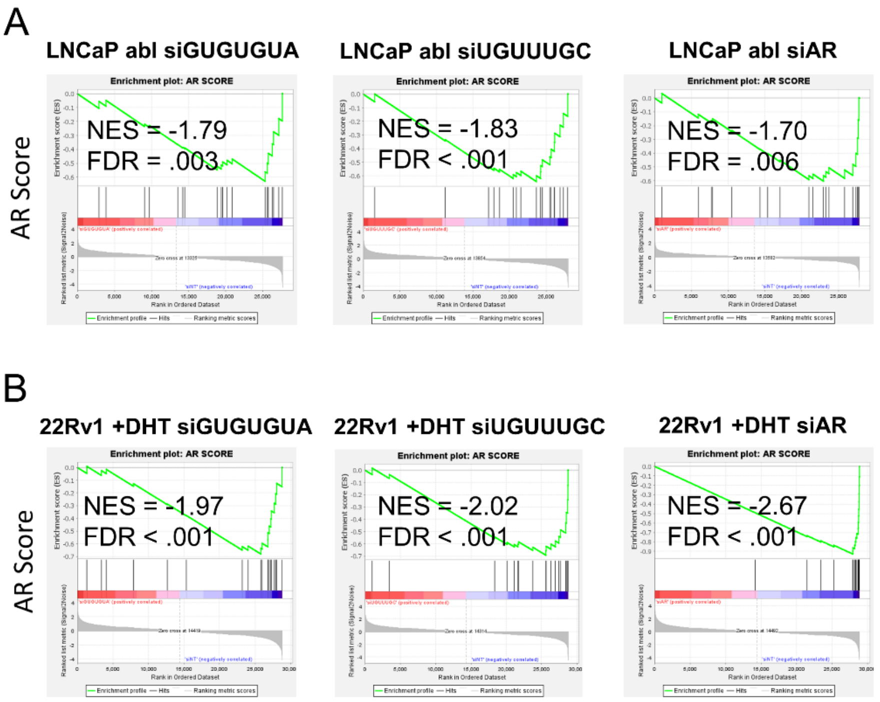


**Figure S16: AR regulated gene sets are downregulated by siGUGUGUA and siUGUUUGC in LNCaP abl and 22Rv1+DHT cells**

GSEA enrichment plots of the “AR score”[30,31] gene set in LNCaP abl (A) and 22Rv1 +DHT (B) cells transfected with siGUGUGUA, siUGUUUGC or siAR.

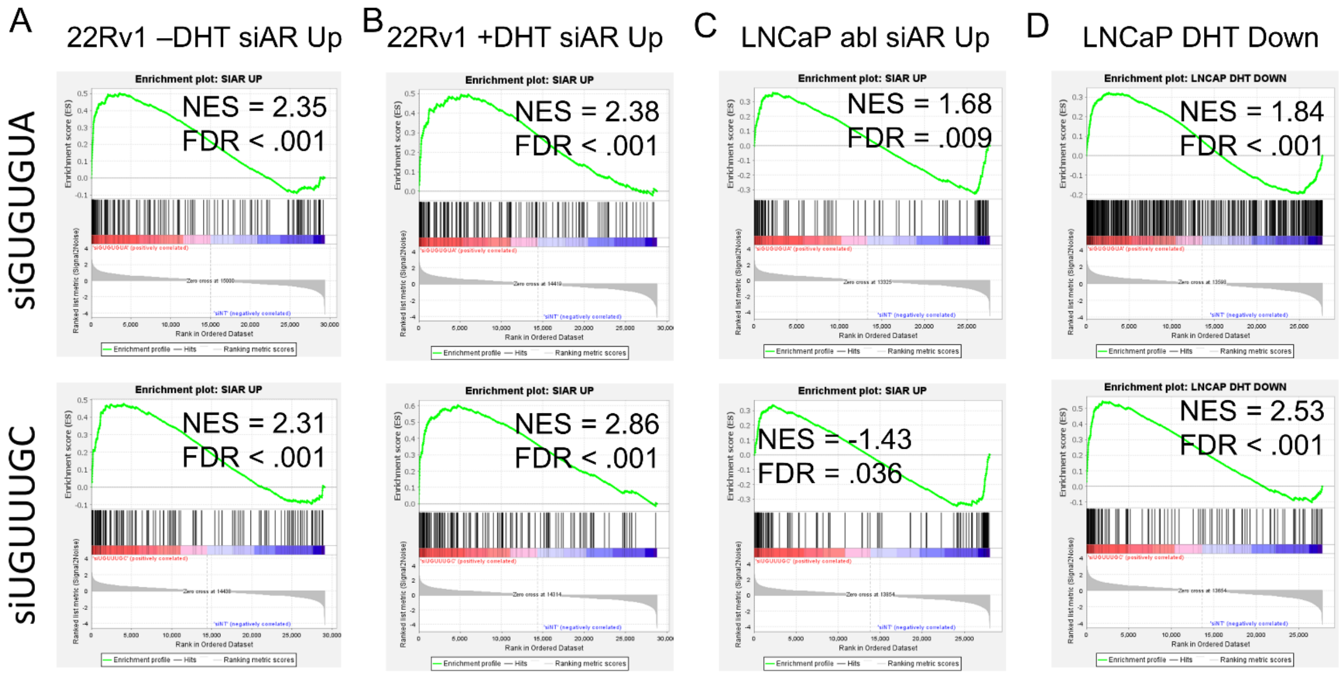


**Figure S17: AR repressed genes exhibit increased expression in response to siGUGUGUA and siUGUUUGC in PCa cells**

GSEA enrichment plots of the "siAR Up" (A-C) and "LNCaP DHT Down"[17] (D) gene sets in 22Rv1 –DHT (A), 22Rv1 +DHT (B), LNCaP abl (C) and LNCaP (D) cells transfected with siGUGUGUA (top) or siUGUUUGC (bottom). Positive and negative NES indicate upregulated and downregulated gene sets respectively. FDR < 0.1 was considered significant.

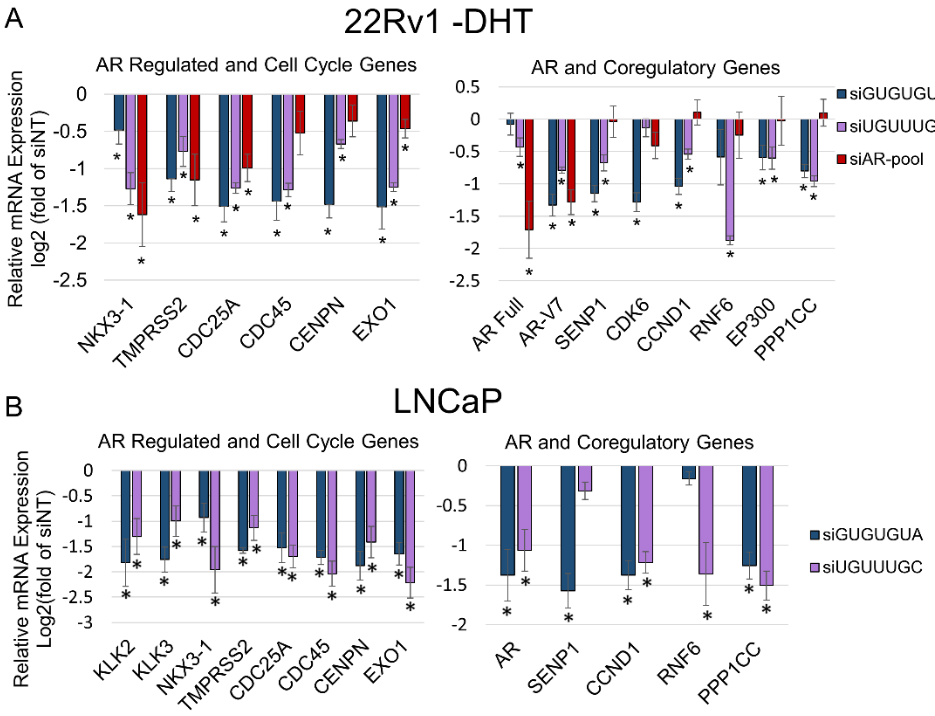


C

| | 3' UTR Length | siGUGUGUA 6mer matches | siGUGUGUA 7mer matches | siUGUUUGC 6mer matches | siUGUUUGC 7mer matches |
|---|---|---|---|---|---|
| KLK2 | 2008 | 2 | 0 | 2 | 1 |
| KLK3 | 637 | 0 | 0 | 0 | 0 |
| NKX3-1 | 2524 | 0 | 0 | 0 | 0 |
| TMPRSS2 | 1837 | 0 | 0 | 0 | 0 |
| CDC25A | 1799 | 0 | 0 | 0 | 0 |
| CDC45 | 129 | 0 | 0 | 0 | 0 |
| CENPN | 2834 | 0 | 0 | 0 | 0 |
| EXO1 | 339 | 0 | 0 | 0 | 0 |
| AR Full | 6778 | 2 | 1 | 5 | 1 |
| AR-V7 | 1353 | 10 | 1 | 1 | 0 |
| SENP1 | 2425 | 4 | 1 | 1 | 0 |
| CDK6 | 10219 | 5 | 2 | 3 | 0 |
| CCND1 | 3192 | 5 | 1 | 1 | 0 |
| RNF6 | 1049 | 0 | 0 | 1 | 1 |
| EP300 | 1121 | 7 | 0 | 2 | 1 |
| PPP1CC | 1396 | 2 | 1 | 1 | 1 |

**Figure S18: AR regulated, cell cycle and AR coregulatory genes are down regulated by siGUGUGUA and siUGUUUGC in LNCaP and 22Rv1-DHT cells**

RT-qPCR validation of downregulated genes identified in RNA seq analysis. Relative mRNA expression of AR regulated, cell cycle, AR coregulators and AR in response to transfection of the indicated siRNA in 22Rv1 -DHT (A) and LNCaP (B) cells. The length of the 3'UTR, and the number of 6mer and 7mer matches to the corresponding siRNA seed in the 3'UTR of each gene are indicated in the table (C). Relative gene expression is expressed as log2(fold) of siNT-1. P < 0.05 was considered significant as determined by t-test, n=3.

A

B

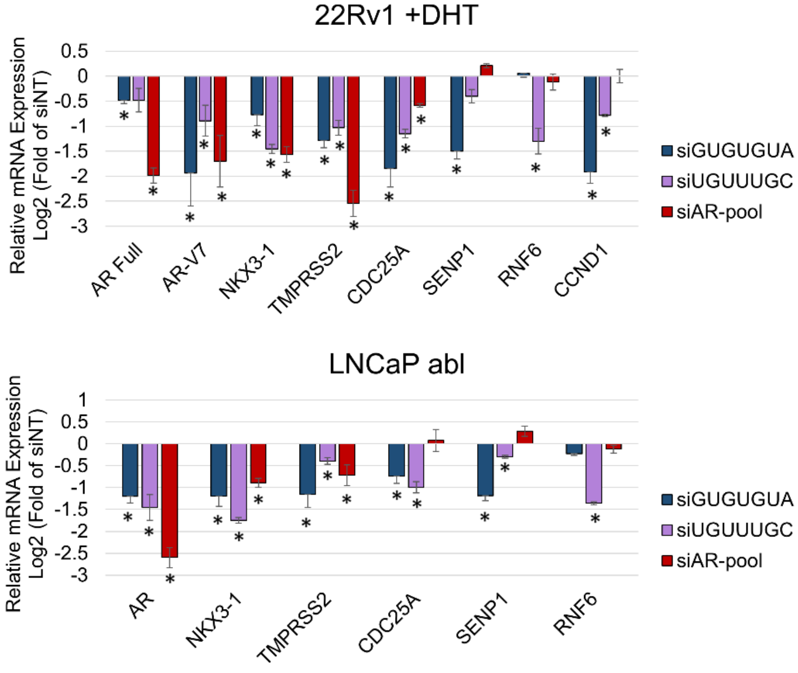


C

| | | siGUGUGUA | | siUGUUUGC | |
|---|---|---|---|---|---|
| | 3' UTR Length | 6mer matches | 7mer matches | 6mer matches | 7mer matches |
| NKX3-1 | 2524 | 0 | 0 | 0 | 0 |
| TMPRSS2 | 1837 | 0 | 0 | 0 | 0 |
| CDC25A | 1799 | 0 | 0 | 0 | 0 |
| AR Full | 6778 | 2 | 1 | 5 | 1 |
| AR-V7 | 1353 | 10 | 1 | 1 | 0 |
| SENP1 | 2425 | 4 | 1 | 1 | 0 |
| CCND1 | 3192 | 5 | 1 | 1 | 0 |
| RNF6 | 1049 | 0 | 0 | 1 | 1 |

**Figure S19: AR regulated, cell cycle and AR coregulatory genes are down regulated by siGUGUGUA and siUGUUUGC in 22Rv1+DHT and LNCaP abl cells**

RT-qPCR validation of downregulated genes identified in RNA seq analysis. Relative mRNA expression of AR regulated, cell cycle, AR coregulators and AR in response to transfection of the indicated siRNA in 22Rv1 +DHT (A) and LNCaP abl (B) cells. The length of the 3'UTR, and the number of 6mer and 7mer seed matches to the corresponding siRNA seed in the 3'UTR of each gene are indicated in the table (C). Relative gene expression is expressed as log2(fold) of siNT-1. $P < 0.05$ was considered significant as determined by t-test, n=3.

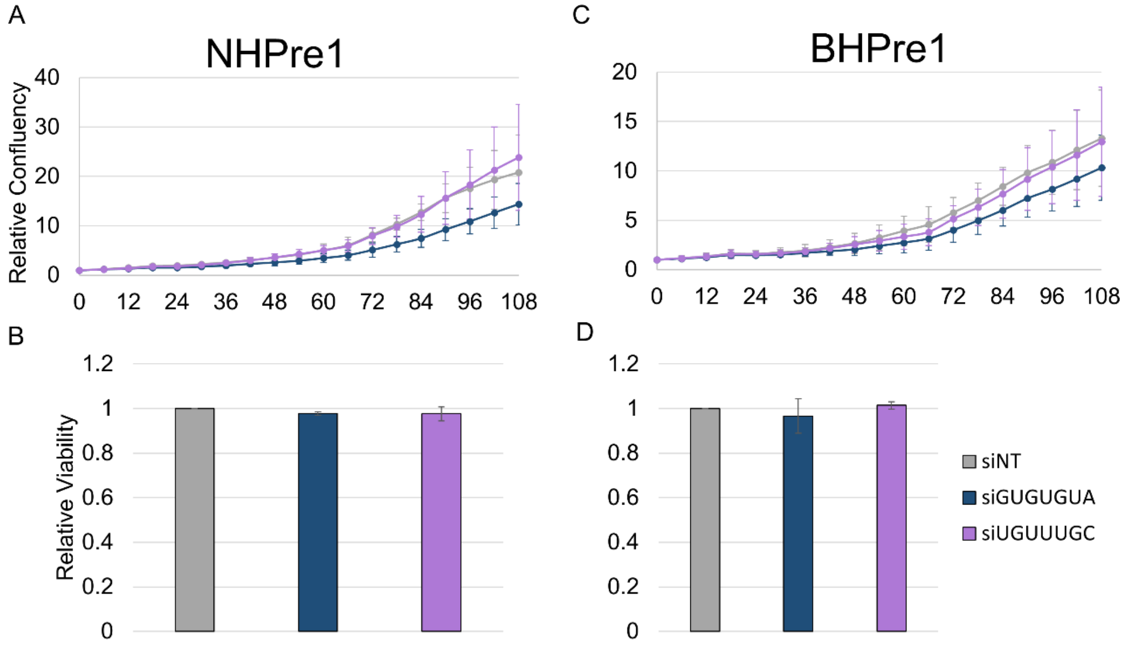


**Figure S20: siGUGUGUA and siUGUUUGC are not toxic in benign prostate cell lines**

Cell growth (A,C) and viability (B,D) of NHPRe1 (A-B) and BHPre1 (C-D) cells transfected with the indicated siRNAs. Relative confluency, as measured by InCucyte, was used as a surrogate for cell growth. Trypan blue was used to measure relative viability. * $p < 0.05$ as determined by T-test, N= 3.

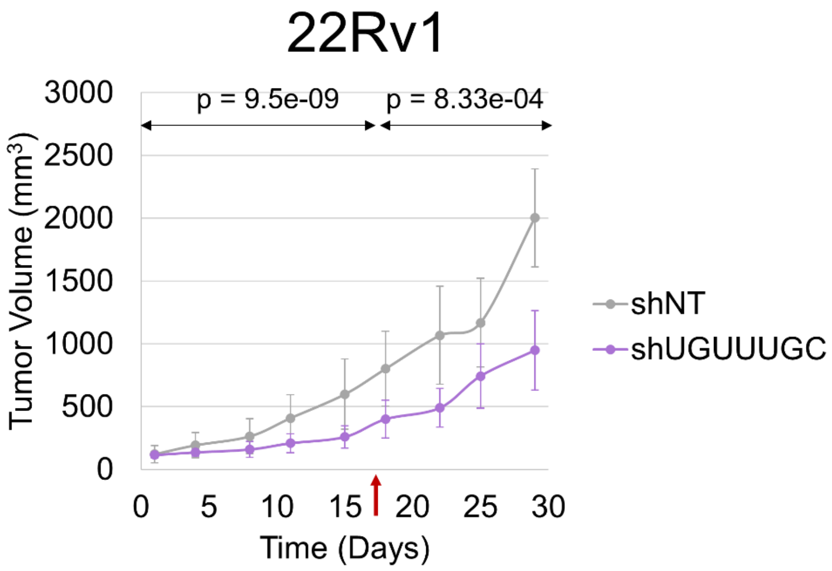


**Figure S21: shUGUUUGC reduces growth of pre-established 22Rv1 xenograft tumors**

Plot showing 22Rv1 shUGUUUGC and shNT tumor growth in mouse xenograft models. Cells were implanted subcutaneously in NOD/SCID mice and fed doxycycline containing chow once tumors were established (volume ≈ 120 $mm^3$). Significance was determined using a segmented linear mixed model, and day 17 was identified as the transition day (red arrow). P values for before and after the transition day are labeled on the plot. n = 6 for shNT; n = 7 for shUGUUUGC.

**Table S1. Number of predicted AR coregulatory and essential genes (LNCaP and general) for the 88 toxic shRNAs**

| Seed | AR Coregulators | | Essential Genes | | LNCaP Essential Genes | |
|---|---|---|---|---|---|---|
| - | # | log10q | # | log10q | # | log10q |
| TATATAT | 59 | 8.487951 | 145 | 2.64843 | 133 | 5.0937961 |
| TATATAC | 43 | 6.0435414 | 98 | 1.384902 | 109 | 5.8648423 |
| CACAGCC | 27 | 6.0435414 | 46 | 0.956394 | 34 | 0.712494 |
| GGCAGTG | 26 | 5.0969387 | 38 | 0.327176 | 46 | 1.975467 |
| GGCAGTA | 20 | 5.0969387 | 24 | 0.319614 | 30 | 1.6339209 |
| TAAGCAG | 21 | 4.6445243 | 36 | 0.792125 | 42 | 3.0256995 |
| TCTGTAC | 24 | 4.5170723 | 59 | 2.56082 | 42 | 1.5856807 |
| TATGTAC | 25 | 4.1976961 | 58 | 1.457608 | 62 | 4.3328984 |
| ACATGGC | 17 | 4.1976961 | 30 | 0.956394 | 21 | 0.6615732 |
| ACAGTAT | 29 | 4.1260354 | 63 | 0.787882 | 65 | 2.4866371 |
| TATGAAC | 21 | 4.1041502 | 46 | 1.410106 | 46 | 3.0256995 |
| GCAACTT | 16 | 3.942317 | 23 | 0.452646 | 34 | 3.1711311 |
| TGTGTGT | 29 | 3.8216488 | 43 | -0.93561 | 63 | 1.8609974 |
| GCAAAGT | 22 | 3.7208483 | 40 | 0.452646 | 50 | 2.7160326 |
| TACACCA | 14 | 3.626331 | 35 | 2.64843 | 27 | 2.3186158 |
| GTGTGTG | 30 | 3.1947372 | 51 | -0.92913 | 65 | 1.1996575 |
| GTGTGTA | 26 | 3.1947372 | 40 | -0.93561 | 58 | 1.6633559 |
| TGGCAGT | 18 | 3.1812655 | 25 | -0.53246 | 37 | 1.7726998 |
| TAAGGAC | 14 | 3.0131001 | 30 | 1.085645 | 41 | 5.3246375 |
| TCCTGTA | 15 | 2.923019 | 33 | 0.992612 | 32 | 2.0114271 |
| TACTGGC | 11 | 2.532899 | 15 | -0.32718 | 24 | 2.1462298 |
| TGGCAGA | 17 | 2.5110411 | 35 | 0.466705 | 35 | 1.2097754 |
| GTGAATG | 14 | 2.5110411 | 24 | 0.332743 | 31 | 1.8758658 |
| TGTGTAA | 15 | 2.4642168 | 37 | 1.048079 | 30 | 1.1996575 |
| TACTGGA | 12 | 2.4642168 | 25 | 0.787882 | 31 | 3.0256995 |
| TATGTTT | 21 | 2.4546956 | 50 | 0.532456 | 54 | 2.111551 |
| TGGGTAA | 11 | 2.4533579 | 25 | 1.085645 | 18 | 0.8554588 |
| GAAACTC | 14 | 2.0377642 | 41 | 1.457608 | 34 | 1.7696502 |
| GCTTGGC | 11 | 2.0377642 | 24 | 0.675825 | 18 | 0.5813691 |
| GACTTGT | 9 | 1.9853208 | 11 | -0.48006 | 18 | 1.3184321 |
| TTAAGGC | 18 | 1.8280968 | 49 | 0.675825 | 53 | 2.4424329 |
| GGCTGGC | 16 | 1.8280968 | 39 | 0.469307 | 52 | 3.5852338 |
| TCCTGGC | 15 | 1.8280968 | 42 | 0.935608 | 34 | 1.0435102 |
| TGTGTAT | 15 | 1.7508518 | 36 | 0.466705 | 36 | 1.2102134 |
| TGTATAC | 13 | 1.7050416 | 39 | 1.215448 | 33 | 1.6097746 |
| TAGGAGT | 8 | 1.6093198 | 16 | 0.469307 | 19 | 1.6318299 |
| CTTACTC | 7 | 1.4923481 | 18 | 0.956394 | 16 | 1.4127431 |
| TAGCATT | 13 | 1.4909744 | 46 | 1.697987 | 38 | 2.0175661 |
| TGTAAGT | 9 | 1.4909744 | 22 | 0.666812 | 25 | 2.0114271 |
| TAAGTAC | 11 | 1.3434034 | 38 | 1.457608 | 31 | 1.7125636 |
| TGGCAGC | 13 | 1.3379502 | 22 | -0.90567 | 37 | 1.6339209 |
| TGGGTAG | 6 | 1.1400083 | 9 | -0.4667 | 11 | 0.5530447 |
| AGAGCAG | 11 | 1.0887495 | 28 | 0.353952 | 29 | 0.9930862 |
| TACTCAG | 8 | 1.0619591 | 22 | 0.584901 | 28 | 2.529876 |
| TCCTGTC | 9 | 1.0523116 | 33 | 1.410106 | 30 | 2.111551 |
| ATGGCTC | 9 | 1.0523116 | 25 | 0.532456 | 23 | 0.9370347 |
| TGCAGGT | 8 | 1.0014805 | 15 | -0.53246 | 17 | 0.456949 |

| | | | | | | |
|---|---|---|---|---|---|---|
| GTGAGAC | 7 | 1.0014805 | 18 | 0.466705 | 16 | 0.6985591 |
| TACTAGG | 5 | 1.0014805 | 14 | 0.781825 | 15 | 1.7130047 |
| TACCTGT | 6 | 0.9677739 | 21 | 1.178987 | 11 | 0.3472926 |
| AGGCAGC | 11 | 0.9499681 | 45 | 1.457608 | 37 | 1.8273787 |
| TGTTTGC | 11 | 0.9414565 | 28 | -0.36241 | 39 | 2.0858776 |
| TAGTGGG | 5 | 0.9103044 | 10 | -0.32718 | 14 | 1.226315 |
| TAAGTCA | 9 | 0.9041347 | 43 | 2.64843 | 43 | 4.7998011 |
| AGGTCTT | 6 | 0.9041347 | 15 | 0.389929 | 16 | 0.9664403 |
| TGTTTAG | 11 | 0.8980656 | 39 | 0.792125 | 49 | 4.0385244 |
| TTCTGGC | 11 | 0.8834726 | 36 | 0.537857 | 49 | 3.9940352 |
| GAATGAG | 11 | 0.86308 | 35 | 0.466705 | 44 | 2.6946382 |
| CCATGGC | 7 | 0.8174871 | 19 | 0.362406 | 22 | 1.2441307 |
| TGCTTGG | 7 | 0.7448986 | 24 | 0.666812 | 23 | 1.2625957 |
| TGTCACT | 7 | 0.7197606 | 26 | 0.792125 | 30 | 2.5587483 |
| CAGAAGC | 12 | 0.6452983 | 45 | 0.554593 | 48 | 2.0114271 |
| GATTCTC | 7 | 0.6452983 | 20 | -0.31961 | 23 | 1.0435102 |
| GCCTGTG | 7 | 0.6318289 | 33 | 1.457608 | 24 | 1.171482 |
| TATAGTG | 5 | 0.6316028 | 11 | -0.46931 | 15 | 0.8038893 |
| TACTCAC | 5 | 0.5899241 | 15 | 0.362406 | 20 | 1.675677 |
| GGTAGCA | 5 | 0.5682859 | 20 | 0.787882 | 19 | 1.3888437 |
| TGATGCC | 4 | 0.5134406 | 21 | 1.457608 | 17 | 1.6339209 |
| CTGTGTG | 9 | 0.4898303 | 39 | 0.771806 | 26 | 0.312553 |
| GATGTGA | 5 | 0.3967252 | 19 | 0.407879 | 19 | 0.872598 |
| ACCTGGC | 4 | 0.3832924 | 18 | 0.666812 | 15 | 0.8018998 |
| GACTGAA | 7 | 0.335419 | 38 | 1.085645 | 40 | 2.8082071 |
| TAGGAGC | 3 | 0.3316442 | 19 | 1.372344 | 18 | 2.111551 |
| TATGAGA | 5 | 0.3280305 | 27 | 0.935608 | 23 | 1.2124911 |
| TCAGCCC | 4 | 0.3280305 | 19 | 0.554593 | 20 | 1.4127431 |
| TACACGC | 1 | 0.3280305 | 3 | 0.327176 | 4 | 0.7740805 |
| TATGTTG | 5 | 0.3274153 | 18 | -0.38936 | 20 | 0.6985591 |
| TTGCAGG | 6 | 0.2921704 | 31 | 0.653227 | 35 | 2.1679562 |
| GACGCCG | 0 | 0 | 1 | 0.365028 | 0 | 0 |
| TCGTCTT | 0 | 0 | 3 | 0.362406 | 3 | 0.5295497 |
| TGGCGGC | 0 | 0 | 3 | 0.466705 | 3 | 0.6985591 |
| ATGAGAC | 4 | -0.3108713 | 24 | 0.823726 | 25 | 1.9765291 |
| GATGCCT | 3 | -0.3274153 | 15 | 0.430595 | 15 | 0.8554588 |
| CGGCAGC | 1 | -0.3280305 | 2 | -0.66681 | 1 | -1.1973459 |
| CATGGCG | 1 | -0.3536195 | 6 | 0.466705 | 6 | 0.7740805 |
| ACTAGGG | 1 | -0.5378029 | 9 | 0.519323 | 8 | 0.722334 |
| TGGTAGT | 1 | -0.9414565 | 17 | 0.773874 | 18 | 1.7279841 |
| TCCTCTA | 3 | -1.0014805 | 30 | 0.430595 | 36 | 2.0033672 |
